# Large scale loss-of-function mutations during chicken evolution and domestication

**DOI:** 10.1101/2024.05.21.595210

**Authors:** Siwen Wu, Kun Wang, Xuehai Ge, Tengfei Dou, Sisi Yuan, Shixiong Yan, Zhiqiang Xu, Yong Liu, Zonghui Jian, Jingying Zhao, Rouhan Zhao, Xiannian Zi, Dahai Gu, Lixian Liu, Qihua Li, Dong-Dong Wu, Junjing Jia, Changrong Ge, Zhengchang Su

## Abstract

Evolution and domestication are often driven by genetic innovations, yet the role of gene loss remains debated. Here, we present comparative genomic analyses of four indigenous chicken breeds from Yunnan Province, China and a reference red jungle fowl genome (GRCg6a). We identify extensive gene presence–absence variation and large numbers of pseudogenes, revealing highly dynamic gene repertoires among closely related chickens. By reconstructing ancestral gene content, we estimate that the common ancestor harbored at least 21,972 genes, of which 7,993 are dispensable. Each lineage has independently lost thousands of genes through both complete gene loss and pseudogenization. These loss-of-function events are non-random: pseudogenization mutations are biased toward gene termini, frequently fixed in populations, and enriched in specific biological pathways. Notably, patterns of gene loss recapitulate phylogenetic relationships, suggesting that loss-of-function mutations are shaped by selection rather than neutral drift. Analysis of four other chicken genomes assembled using PacBio HiFi reads draws the same conclusion. Thus, our results support a model in which large-scale loss-of-function mutations are a major driver of chicken evolution and domestication, consistent with the “less-is-more” hypothesis of adaptive evolution.

## Introduction

As the first non-mammalian vertebrate sequenced [1], chicken (*Gallus gallus*) provides us with most protein sources in our daily life and also is an important model organism to study development and immunity of vertebrates [2]. Since the first release of the draft genome of a red jungle fowl (RJF) [1], its assembly quality has been greatly improved (galgal5 and GRCg6a) [3, 4]. Moreover, the Vertebrate Genomes Project (VGP) assembled pseudo-haplotype genomes (GRCg7b and GRCg7w) of a hybrid individual from a broiler mother and a layer father using long sequencing reads and multiple types of scaffolding data [5, 6]. More recently, genomes of several chicken breeds such as Huxu [7], Hailanhe [8], Silkie [9] and Lueyang [10] were assembled using PacBio HiFi reads with improved quality. Studies based on these assemblies have provided insightful understandings of not only the domestication and evolution processes, but also the genetic basis of selected traits of domesticated chickens. However, these earlier studies were limited in revealing driving forces of chicken domestication and evolution due to either low-quality genome assemblies or insufficient gene annotations. Consequently, contradictory conclusions have been drawn. For example, on the one hand, it was reported that the chicken genome has undergone a large number of segmental deletions [1], resulting in a substantial reduction of genome sizes and a large number of concomitant gene loss; and therefore, chicken might have fewer genes than other tetrapods [11]. On the other hand, it was concluded that selection for loss-of-function mutations had no prominent role in chicken domestication [12, 13]. However, accumulating genomic evidence supports loss-of-function mutations as a major driving force for the evolution (for a review, see [14–16]) of animals [17–21] and plants [22] as well as for the domestication (for a review, see [23, 24]) of many farm animals [25] and crops [26, 27].

Furthermore, it has been shown that RJF subspecies *G. g. Spadiceus* is the primary ancestor of domestic chickens (*Gallus gallus domesticus*) all over the world [28]. *G. g. Spadiceus* diverged from other RJF subspecies *G. g. murphy*, *G. g. jabouillei*, *G. g. gallus* and *G. g. bankiva* 50,000-500,000 years ago [28], substantially earlier than the advent of chicken domestication [29]. Indigenous chickens in Yunnan, a southwestern province in China, are among the earliest domesticated birds, and they are formed by less-intensive traditional family-based artificial selection in villages in isolated mountainous areas since 2,000–6,000 BC [29]. It has been estimated that indigenous chickens in Yunnan share the most recent common ancestor (MRCA) with wild *G. g. Spadiceus* less than 8,000 years ago [28]. Although domestic indigenous chicken such as those from southeast Asia and commercial chickens such as white leghorn may have substantial introgression from other RJF subspecies, indigenous chickens in Yunnan have minimal (∼4%) introgression only from *G. g. jabouille* [28]. Thus, indigenous chickens in Yunnan are good candidates to investigate the driving force of evolution and domestication of indigenous chickens.

We therefore recently sequenced and assembled genomes at chromosome-level of four indigenous chicken breeds in Yunnan [30]. These chicken breeds include Daweishan chicken with a miniature body size, Hu chicken with a large body size and stout legs, Piao chicken with a rumpless trait, and Wuding chicken good at running. Using an annotation pipeline that combines homology-based and RNA-seq-based methods, we found that the Daweishan, Hu, Piao and Wuding chicken genomes encoded 17,718, 17,497, 17,711 and 17,646 protein-coding genes, respectively [31]. Of these genes in the four genomes, a total of 1,420 are not seen in the annotations of the RJF (GRCg6a), the broiler (GRCg7b) and the layer (GRCg7w) assemblies, we thus refer them to as newly annotated genes (NAGs) [31]. Unexpectedly, we also identified a large number of pseudogenes in the Daweishan (747), Hu (606), Piao (682) and Wuding (667) chicken genomes [31]. Interestingly, most of the NAGs also are either encoded or become pseudogenes in the GRCg6a, GRCg7b, and GRCga7w assemblies. We therefore increase the numbers of both annotated genes and pseudogenes in GRCg6a (18,463 and 542), GRCg7b (19,002 and 474) and GRCg7w (18,978 and 435) [31]. In addition to the varying numbers of genes and pseudogenes in these genomes, each pair of breeds share only 81%-92% of their genes. This is in stark contrast to the observation that individual genomes of some species such as humans encode almost the same set of protein-coding genes, regardless of their ancestral origin [32–35]. Moreover, even some closely-related different species such as humans and chimpanzees share 98% of their genes [36]. To understand the underlying reasons for such high variation in gene and pseudogene compositions in the chickens, we analyzed the occurring patterns and evolutionary behaviors of the pseudogenes as well as the presence and absence variation of genes in the four highly closely-related indigenous chicken genomes together with the RJF genome. Our results suggest that loss-of-function of genes via a pseudogenization mutation (a nonsense mutation resulting in a premature stop codon or an insertion/deletion mutation leading to an open-reading-frame (ORF) shift) or complete gene losses (a gene cannot be detected even as a pseudogene) might play critical roles in chicken evolution and domestication. This conclusion is also supported by the results from analyzing the Huxu chicken [7], Hailanhe chicken [8], Silkie chicken [9] and Lueyang chicken [10] genomes assembled using PacBio HiFi reads.

## Results

### Chicken genomes harbor highly varying sets of protein-coding genes and pseudogenes

Although the Daweishan (17,718), Hu (17,497), Piao (17,711) and Wuding (17,646) chicken genomes harbor quite similar numbers of protein-coding genes [31], they shared only 15,050 genes (Figure 1a), comprising only 84.9%-86.0% of their genes. Interestingly, the indigenous chicken genomes encode 745-966 fewer genes than the RJF genome (GRCg6a) (18,463). Consequently, the indigenous chicken genomes share only 13,979 genes with the RJF genome (Figure 1b), comprising 75.7%-79.9% of their genes. Moreover, each pair of the four indigenous chicken genomes share only 91%-92% of their genes (Figure 1c), even though they diverged only 8,000-years ago [28]. In contrast, different ancestral groups of some mammal species such as humans share almost the same set of protein-coding genes [32–35], and even some closely related mammal species such as humans and chimpanzees share 98% of their genes although they split 6-7 million years ago [36].

**Figure 1.**
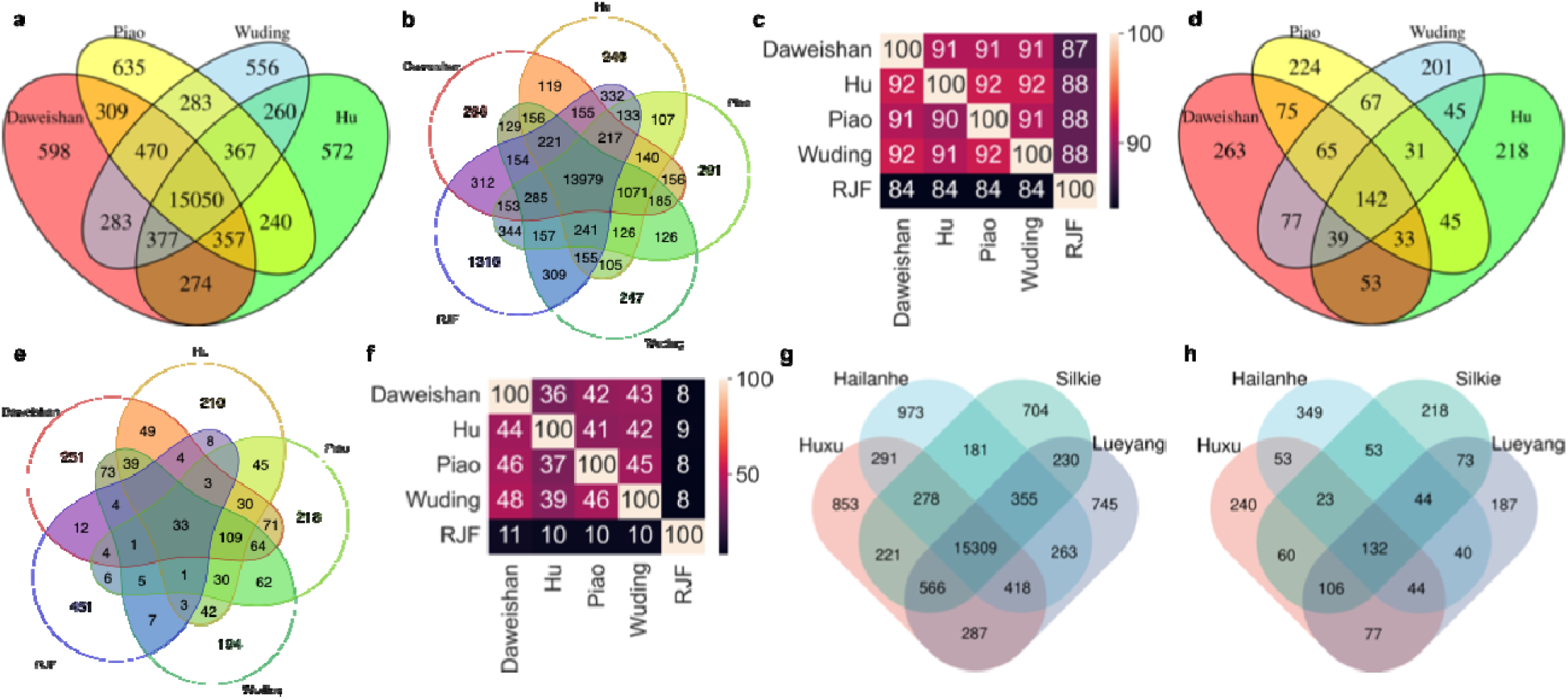
Comparison of protein-coding genes and pseudogenes among different chicken breeds. **a.** Venn diagram of the protein-coding genes of our four indigenous chickens. **b.** Venn diagram of the protein-coding genes of our four indigenous chickens and RJF. **c.** Comparison of the protein-coding genes among each pair of our four indigenous chickens and RJF. **d.** Venn diagram of the pseudogenes of our four indigenous chickens. **e.** Venn diagram of the pseudogenes of our four indigenous chickens and RJF. **f.** Comparison of the pseudogenes among each pair of our four indigenous chickens and RJF. **g.** Venn diagram of the protein-coding genes of the four HiFi-reads assembled genomes. **h.** Venn diagram of the pseudogenes of the four HiFi-reads assembled genomes.

Moreover, we identified a larger number of pseudogenes in each of the four indigenous chicken genomes (606-747) (Table 1) based on three sources [31] (Materials and Methods). Most (486-622 or 80.2%-83.5%) of the pseudogenes in the indigenous chicken genomes were predicted based on homology to annotated genes in GRCg6a (322-395), GRCg7b (313-385) and GRCg7w (311-398) (Tables 1 and S1-S4). In other words, some functional genes in the reference genomes become pseudogenes in an indigenous chicken genome due at least to one pseudogenization mutation. A small portion (24-35 or 3.6%-5.8%) were predicted based on homology to the 1,420 NAGs [31], i.e., some functional NAGs in indigenous genomes become pseudogenes in other indigenous genomes. The remaining 83-94 (12.2%-14.0%) were predicted based on homology to previously annotated pseudogenes in GRCg6a (49-57), GRCg7b (55-64) and GRCg7w (51-58). In other words, some pseudogenes in reference genomes also are pseudogenes in indigenous chicken genomes. Most pseudogenes (576-713, 94.9-96.0%) in each indigenous chicken genome are transcribed in multiple tissues (Tables S5-S8), suggesting that their regulatory systems might be still at least partially functional. Furthermore, based on pseudogenization mutations of the 1,420 NAGs in the reference genomes [31], we increased the numbers of annotated pseudogenes in GRCg6a (from 262 to 542) (Table 1). Notably, the indigenous chicken genomes harbor 64-205 more pseudogenes than GRCg6a, which partially explains why the former genomes harbor fewer genes than the latter genome (17,497-17,716 vs 18,463). The indigenous chickens share 142 pseudogenes (Figure 1d) among themselves, and 33 pseudogenes with the RJF (Figure 1e). Each pair of the chicken share 8%-48% of their pseudogenes (Figure 1f). In total, we found 1,995 pseudogenes that appeared in at least one of the five chicken genomes (Table S9).

**Table 1:**
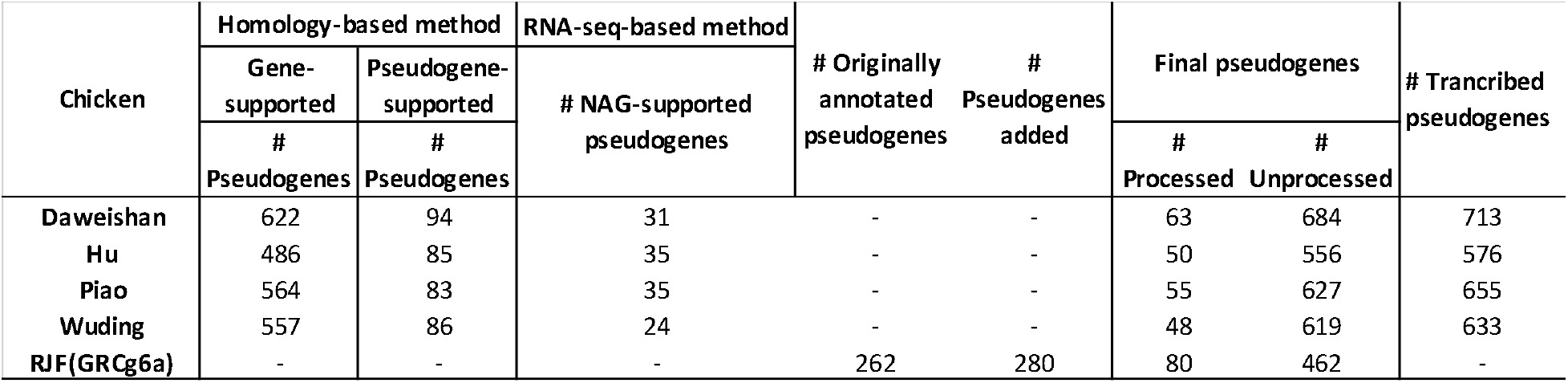
Summary of annotated pseudogenes in the four indigenous chicken genomes in comparison with those in GRCg6a.

To see whether or not such large variation in the numbers of annotated protein-coding genes as well as the large numbers of pseudogenes in our assembled genomes and the GRCg6a assembly is due to their assembly quality, as they were assembled using a combination of error-prone Nanopore/PacBio long reads and highly accurate Illumina short reads [30], we evaluated and annotated (Materials and Methods) PacBio HiFi-reads assembled genomes of four other chicken breeds [7–10], including a Huxu chicken (GGswu), a Silkie chicken (ASM3450986v1), a Lueyang chicken (ASM3450988v1) and a Hailanhe chicken (ASM3450984v1) (Table 2). Huxu, Silkie and Lueyang are indigenous breeds in China, while Hailanhe is a layer derived from a breed from Hy-Line International. We found that all the four HiFi-reads assemblies had a BUSCO completeness value of 99% based on the BUSCO (5.7.1) aves_odb10 database [37], suggesting that they are indeed of higher quality than our four indigenous chicken assemblies and GRCg6a with a BUSCO completeness value of ∼96% [30]. Using the coding sequences (CDSs) of GRCg6a, GRCg7b and GRCg7w assemblies as well as our 1,420 NAGs [31] as the references, we annotated from 17,844 (Silkie) to 18,223 (Huxu) protein-coding genes and from 703 (Lueyang) to 735 (Huxu) pseudogenes in each of the four assemblies (Tables 2 and S10-S13). Thus, we were able to annotate more protein-coding genes (17,844-18,223 vs. 17,497-17,718) as well as more pseudogenes (703-735 vs. 606-704) in these HiFi-reads assemblies due probably to their more complete assembly than in our assembled indigenous genomes. Nonetheless, like our assembled indigenous genomes, these HiFi-reads assemblies also shared only 15,309 genes (Figure 1g), comprising only 84.0%-85.6% of their genes. Like our assemblies, these HiFi-reads assemblies also encode 240-619 fewer genes than the RJF genome (GRCg6a) (18,463) that was assembled using a combination of error-prone Nanopore/PacBio long reads and highly accurate Illumina short reads [30]. Moreover, like our assemblies, these HiFi-reads assemblies also harbor 161-193 more pseudogenes than GRCg6a (542), which also partially explains why the former genomes harbor fewer genes than the latter genome (17,844-18,223 vs. 18,463). Furthermore, like our assemblies, these HiFi-reads assemblies share 132 pseudogenes (Figure 1h), comprising only 18.0%-18.9% of their pseudogenes. Therefore, the high variation in the number of annotated genes as well as the large numbers of annotated pseudogenes in our assemblies are unlikely due mainly to their relatively lower assembly quality.

**Table 2:** Summary of annotated protein-coding genes, pseudogenes and substitution rates in the four PacBio HiFi-reads assembled chicken genomes.

| Assembly | Breeds | # Protein-coding genes | # Pseudogenes |  |  | Mean substitution rates of protein-coding genes | Mean substitution rates of pseudogenes |
| --- | --- | --- | --- | --- | --- | --- | --- |
|  |  |  | # Unprocessed | # Processed | # Unprocessed with duplication |  |  |
| GGswu | Huxu | 18223 | 706 | 29 | 106 | 4.87e-09 | 1.03e-08 |
| ASM3450984v1 | Hailanhe | 18068 | 687 | 51 | 122 | 4.47e-09 | 1.33e-08 |
| ASM3450986v1 | Silkie | 17844 | 659 | 50 | 112 | 4.86e-09 | 1.75e-08 |
| ASM3450988v1 | Lueyang | 18173 | 667 | 36 | 122 | 4.69e-09 | 1.62e-08 |

### Vast majority of pseudogenes are unprocessed and unitary

Most (91.6%-92.8%, 556-684) of the pseudogenes in each of our indigenous chicken genome are unprocessed (Table 1), i.e., they arose due to direct pseudogenization mutations, while the remaining small portion (7.2-8.4%, 48-63) are processed, i.e., they arose due to retrotransposition followed by pseudogenization mutations [38]. Unprocessed pseudogenization mutations could occur after a duplication event to eliminate a redundant copy [38]. However, we failed to find an intact paralog for most (88.2%-90.3%) of the unprocessed pseudogenes in the same genomes (Tables S1-S4), suggesting that most of the unprocessed pseudogenes are not related to gene duplications, and thus are unitary pseudogenes [39]. There are a total of 1,814 unprocessed pseudogenes in our four chickens and the RJF genomes (Table S14). The indigenous chickens share 22 unprocessed pseudogenes with the RJF (Figure S1a) and 129 among themselves (Figure S1b). Interestingly, compared to the cases in indigenous chickens, a smaller proportion of the pseudogenes in RJF (85.2%, 462) are unprocessed. However, the number (80) of processed pseudogenes in RJF is similar to those (63, 50, 55 and 48) in the four indigenous chickens (Table 1). Similarly, most (659-706) of the predicted pseudogenes (703-735) in the four HiFi-reads assemblies are also unprocessed, and only 29-51 were processed (Tables 2 and S10-S13). Moreover, among these unprocessed pseudogenes, only a few (106-122) of them have functional copies in their genomes (Tables 2 and S10-S13).

### Pseudogenization mutations are biased to the two ends of parental genes

When a pseudogene may contain more than one pseudogenization mutation, the most upstream one (defined as the one that is closest to the 5’-end of the parental gene) obviously has the highest impact on the sequence of the pseudogene. Thus, to see whether the arise of the unprocessed pseudogenes in the indigenous chickens is under natural/artificial selection or selectively neutral, we compared the distribution of the relative positions of their most upstream pseudogenization mutations along the CDSs of their parental genes with the distribution of the relative positions of synonymous mutations along the CDSs of true genes. Synonymous mutations in true genes in both of our assembled genomes (Figure 2a) and HiFi-reads assembled genomes (Figure 2b) are largely uniformly distributed along the CDSs as expected for neutral mutations, except at the two ends, where the substitution rate decreases, consistent with an earlier report in chickens [40]. The reduced synonymous substitution rates at the two ends of true genes suggest that the two ends of CDSs are generally under stronger purifying selection and that the two ends might harbor functional elements not related to their amino acid coding functions, such as transcriptional and post-transcriptional regulatory elements [41]. Interestingly, the synonymous mutations in the pseudogenes in both our four indigenous chickens and the HiFi-reads assembled genomes are also largely uniformly distributed along the CDSs, except at the two ends (Figures 2a and 2b). However, the extent to which synonymous substitution rates reduced at the two ends of pseudogenes is smaller than that for the true genes (Figures 2a and 2b), indicating that purifying selection on the two ends of pseudogenes is relaxed. Thus, transcriptional and post-transcriptional regulatory elements at the two ends of pseudogenes might have been deteriorated since their pseudogenization.

**Figure 2.**
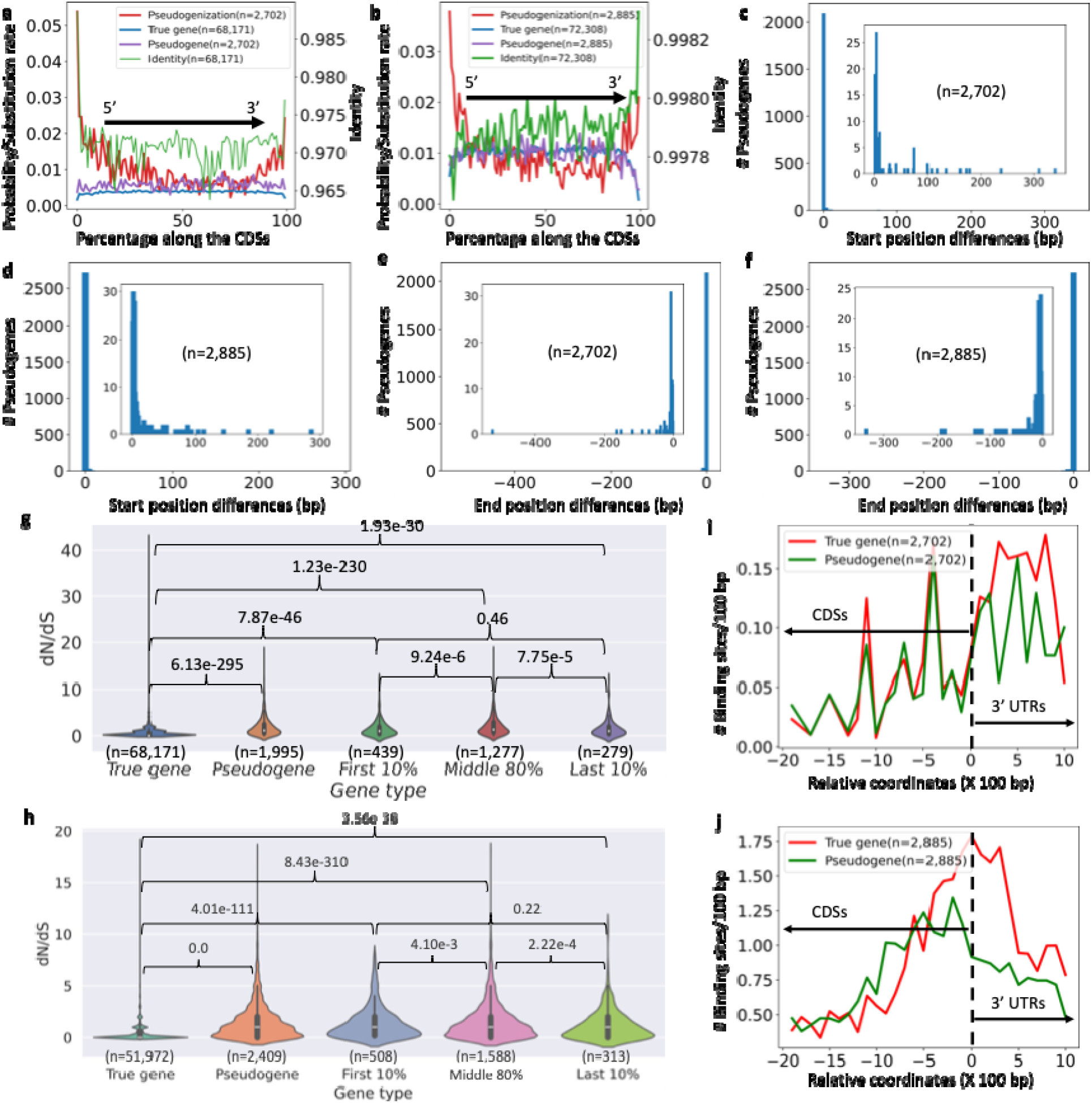
Pseudogenization mutations tend to occur at the two ends of CDSs. **a-b.** Probability of the most upstream pseudogenization mutations (red line) in 100 evenly divided CDS segments from the 5’-ends to the 3’-ends of the parental genes of the pseudogenes, mean rates of synonymous substitution in 100 evenly divided CDS segments from the 5’-ends to the 3’-ends of the true genes (blue line) and pseudogenes (purple), and mean identity of the true genes in 100 evenly divided CDS segments from the 5’-ends to the 3’-ends of the genes (green line). (a: For our four indigenous chickens; b: For the four HiFi-reads assembled genomes). **c-d.** Start position of the “CDS” of the pseudogenes with respect to the nucleotide positions of their parental genes starting with 0 with the downstream positions being positive integers (c: For our four indigenous chickens; d: For the four HiFi-reads assembled genomes). **e-f.** End positions of the “CDS” of the pseudogenes with respect to the nucleotide positions of their parental genes ending with 0 with the upstream positions being negative integers (e: For our four indigenous chickens; f: For the four HiFi-reads assembled genomes). **g-h.** Violin plots of the dN/dS values of all true genes, all pseudogenes, pseudogenes with the most upstream pseudogenization occurring in the first 10%, the intermediate 80% and the last 10% of the CDSs (g: For our four indigenous chickens; h: For the four HiFi-reads assembled genomes). **i-j.** Number of predicted miRNA binding sites per 100pb along the CDSs and 3’ UTRs of the true genes (red line) and pseudogenes (green line). In the figure, ‘0’ represents the end positions of the CDSs, the positive numbers represent the relative positions of 1,000 bp sequences downstream of the end of CDSs, and the negative numbers represent the relative positions of the CDSs with respect to the ends of CDSs (i: For our four indigenous chickens; j: For the four HiFi-reads assembled genomes).

By stark contrast, the most upstream pseudogenization mutations in the pseudogenes in both our assembled genomes (Figure 2a) and the HiFi-reads assembled genomes (Figure 2b) are strongly biased to the 5’-ends and 3’-ends of the parental CDSs, consistent with earlier reports in chickens [40] and humans [42]. Specifically, 22.2%, 64.2% and 13.6% of the most upstream pseudogenization mutations in our assembled genomes occur in the first 10%, the middle 80% and the last 10% lengths of the CDSs (counting from the 5’-end to the 3’-end of the CDSs) (Figure 2a), and similar results (21.4%, 64.9% and 13.7%, respectively) are obtained in the four HiFi-reads assembled genomes (Figure 2b). Almost all the pseudogenes in both our assembled genomes and the HiFi-reads assembled genomes have their first (Figures 2c and 2d) and last (Figures 2e and 2f) coding nucleotides aligned with those of parental CDSs, indicating that the biased pseudogenization mutations to the 5’- and 3’-ends are not due to incorrect predictions of the two ends of the pseudogenes. These results strongly suggest that the biased most upstream pseudogenization mutations towards the two ends of pseudogenes are under positive selection or artificial selection.

### The most upstream pseudogenization mutations biased to the two ends of parental CDSs might facilitate loss of functions of genes

To see whether the most upstream pseudogenization mutations along the parental CDSs result in loss of function of the genes, we compared the evolutionary pressure on true genes with that on pseudogenes using the ratios of the number of nonsynonymous mutations over the number of synonymous mutations (dN/dS). The pseudogenes have significantly higher dN/dS values than the true genes in both our assembled genomes (p=6.13e-295, Figure 2g) and the HiFi-reads assembled genomes (p=1e-325, Figure 2h). This is also true when the pseudogenes with the most upstream pseudogenization mutations occurring in the first 10% (p=7.87e-46, Figure 2g; p=4.01e-11, Figure 2h), the middle 80% (p=1.23e-230, Figure 2g; p=8.43e-310, Figure 2h) or the last 10% (p=1.93e-30, Figure 2g; p=3.56e-38, Figure 2h) lengths of parental CDSs are compared with the true genes (Wilcoxon rank-sum test). These results suggest that at least most pseudogenes are no longer under purifying selection, and thus might lose gene functions. Moreover, the pseudogenes with the most upstream pseudogenization mutations occurring in the first 10% lengths of parental CDSs and occurring in the last 10% lengths of parental CDSs have similar dN/dS values (p=0.46, Figure 2g; p=0.22, Figure 2h), which are slightly but significantly lower than those of the pseudogenes with the most upstream pseudogenization mutation occurring in the middle 80% lengths of parental CDSs in both our assembled genomes (p=9.24e-6 and p=7.75e-5, respectively, Wilcoxon rank-sum test, Figure 2g) and the HiFi-reads assembled genomes (p=4.10e-3 and p=2.22e-4, respectively, Wilcoxon rank-sum test, Figure 2h). The underlying cause is not clear to us but might be due to our finding that the two ends of CDSs were under stronger purifying selection before pseudogenization events occurred (Figures 2a and 2b).

Clearly, the closer a pseudogenization mutation to the 5’-end of the CDSs, the larger portion of the peptide chain is affected, and the more likely a pseudogenization mutation occurs. Loss of function of a gene could also occur when critical amino acids at the C-terminus of the protein or regulatory DNA elements at the 3’-end of the CDSs are disrupted by a pseudogenization mutation. For the former possibility, we noted that the identity of amino acids at the C-terminus of proteins are elevated in both our assembled genomes (Figure 2a) and HiFi-reads assembled genomes (Figure 2b), indicating that the C-terminus may harbor critical amino acids whose mutations can significantly affect protein functions. For the latter possibility, as the 3’-UTRs of genes often harbor miRNA binding sites for post-transcriptional regulation [43], we hypothesize that 3’-ends of CDSs may also contain miRNA binding sites, and disruption of such sites in either 3’-ends of CDSs or 3’-UTRs may have functional consequence. To test this, we scanned the pseudogenes’ and their parental genes’ CDSs and their 1,000 bp downstream sequences as putative 3’-UTRs for potential miRNA binding sites. Putative 3’-UTRs and 3’-ends of both parental genes and pseudogenes in both our assembled genomes (Figure 2i) and the HiFi-reads assembled genomes (Figure 2j) have higher density of putative miRNA binding sites than their upstream coding regions, consistent with previously reports [43], suggesting that putative 3’-UTRs and 3’-ends indeed tend to contain miRNA binding sites. Interestingly, pseudogenes have a fewer number of miRNA binding sites in their 3’-ends and 3’-UTRs than do parental true genes (Figures 2i and 2j), suggesting that pseudogenization mutations might disrupt the miRNA binding sites in the 3’-ends of CDSs.

### Vast majority of the most upstream pseudogenization mutations are fixed in respective populations

To see whether the most upstream pseudogenization mutations along a pseudogene in our four indigenous chicken genomes are fixed or not in their respective populations, we computed the frequencies of the mutated alleles in the populations of the four breeds based on DNA re-sequencing reads (Materials and methods). As shown in Figures 3a-3d, most (∼75%) of the most upstream pseudogenization mutations in the pseudogenes in each chicken genome are fixed or nearly fixed (allele frequency > 80%) in their respective populations. The high probability of fixation of the most upstream pseudogenization mutations suggests again that such mutations might be under positive selection to eliminate the functions of the parental genes. A few examples of fixed or nearly fixed unitary pseudogenes in indigenous chicken populations are shown in Figure S2. Since population data are not available for most of the four HiFi-reads assemblies, we did not analyze the fixation rate of their pseudogenes.

**Figure 3.**
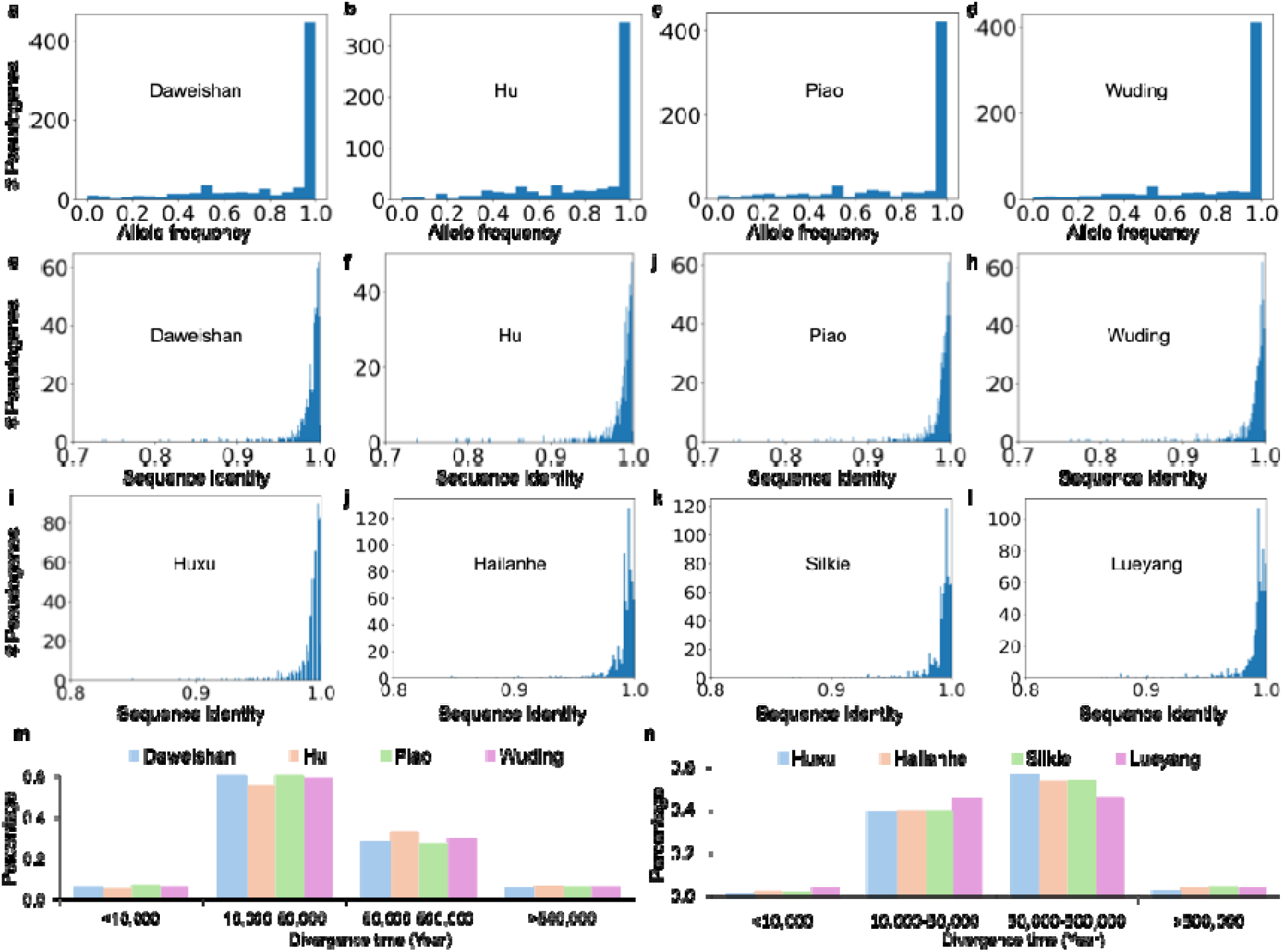
Spectrum analysis of pseudogenes in our four chicken breeds and the four HiFi-reads assemblies. **a-d.** Number of pseudogenes with the indicated fixation rate of the most upstream pseudogenization mutations in the chicken populations. **e-h.** Number of pseudogenes in each of our indigenous chicken genomes with the indicated identity with their parental genes. **i-l.** Number of pseudogenes in each of the four HiFi-reads assembled genomes with the indicated identity with their parental genes. **m-n.** Distribution of divergence times of pseudogenes in each chicken breed since the divergence from their last common ancestral genes shared with the RJF (GRCg6a).

### The functions of parental genes of most pseudogenes are lost in our indigenous chicken genomes

To see whether alternative isoforms of the pseudogenes in the four indigenous chickens could skip the exons harboring the pseudogenization mutations, we assembled transcripts of all the transcribed pseudogenes in each of our indigenous chicken genomes (Table 1). We found that most transcribed pseudogenes had only one type of transcript containing the pseudogenization mutations, while for those that had more than one isoform, very few of them had transcripts that skipped the pseudogenization mutations (Tables S15-S18). For example, in Daweishan chicken, only 139 (20.32%) of the 684 unprocessed pseudogenes (Table 1) have alternative splicing transcripts, and only one of them has transcripts that skip the pseudogenization mutation (Table S15). In Hu chicken, only 137 (24.64%) of the 556 unprocessed pseudogenes (Table 1) have alternative splicing transcripts, and none of them has transcripts that skip the pseudogenization mutations (Table S16). In Piao chicken, only 131 (20.89%) of the 556 unprocessed pseudogenes (Table 1) have alternative splicing transcripts, and only two of them have transcripts that skip the pseudogenization mutations (Table S17). In Wuding chicken, only 139 (22.46%) of the 619 unprocessed pseudogenes (Table 1) have alternative splicing transcripts, and none of them has transcripts that skip the pseudogenization mutations (Table S18). These results suggest that almost all the pseudogenes in the four indigenous chickens did not skip the exons harboring the pseudogenization mutations, and that the functions of parental genes cannot be rescued by alternative isoforms of the pseudogenes. Moreover, as we indicted earlier, most of the unprocessed pseudogenes do not have a functional copy in the same genomes, thus, the functions of their parental genes might be lost in the indigenous chickens. These findings suggest again that most of the pseudogenes in our four indigenous chickens have lost their functions. However, the lack of RNA-seq data in the four PacBio HiFi-assemblies prevented us from conducting a similar analysis.

### Most pseudogenes arose during the subspeciation and domestication processes

The vast majority (93%-97%) of the pseudogenes in our four indigenous chicken genomes (Figures 3e-3h) as well as in the four HiFi-reads assembled genomes (Figures 3i-3l) share more than 95% of sequence identity with their parental genes, indicating that they arose quietly recently, while only a small portion (0%-1%) with less than 80% of sequence identity with their functional parental genes arose a relatively long time ago. To estimate the ages of the pseudogenes, we employed the Kimura’s two-parameter (K2P) model [44] and computed the mean substitution rates (λ) (number of fixations per year per site) of the pseudogenes/protein-coding genes in each chicken breed with respect to the ortholog sequences in the genome of the RJF (GRCg6a) (Materials and Methods). The pseudogenes in each breed with orthologous sequences identified in the GRCg6a assembly are listed in Tables S19-S26. As expected, pseudogenes have several folds higher mean substitution rates than do protein-coding genes in each of our assembled genomes (1.04e-08 vs 1.16-e09 for Daweishan, 1.12e-08 vs 1.41e-09 for Hu, 1.02e-08 vs 1.10e-09 for Piao, and 1.08e-08 vs 1.10e-09 for Wuding, Table 3) as well as in each of the HiFi-reads assembled genomes (1.03e-08 vs 4.87e-09 for Huxu, 1.33e-08 vs 4.47e-09 for Hailanhe, 1.75e-08 vs 4.86e-09 for Silkie, and 1.62e-08 vs 4.69e-09 for Lueyang, Table 2), presumably due to stronger purifying selection on the protein-coding genes than on pseudogenes (Figures 2g and 2h). Based on the calculated mean substitution rate λ, we estimated the divergence time (age) of each pseudogene in each breed with the orthologues genes in the RJF (GRCg6a) (Tables S19-S26). As summarized in Figure 3m for our assembled genomes, 5.2%-6.5% pseudogenes appeared in the past 10,000 years, 55.8%-60.2% arose in 10,000-50,000 years ago, 27.3%-32.9% emerged 50,000-500,000 years ago, and the remaining 5.6%-6.1% occurred more than 500,000 years ago. Similar results were obtained for the HiFi-reads assembled genomes (Figure 3n). Specifically, 1.4%-3.8% pseudogenes appeared in the past 10,000 years, 39.2%-45.8% arose in 10,000-50,000 years ago, 46.4%-56.6% emerged 50,000-500,000 years ago, and the remaining 2.9%-4.4% occurred more than 500,000 years ago. It has been reported that the domestication of chickens in southwest China including Yunnan [29] as well as in north China [45] began 10,000 years ago. Thus, 1.4%-6.5% pseudogenes that appeared during this period of time might be resulted from a combination of artificial selection and natural selection. Moreover, it has been estimated that the subspeciation of *G. gallus* started 50,000-500,000 years ago [28]. Hence, 27.3%-56.6% pseudogenes might arise during this period of time, while 39.2%-60.2% might emerge in the evolution process after subspeciation. These results suggest that most (93.9%-97.1%) pseudogenes might arose during the processes of subspeciation and domestication.

**Table 3:** Mean substitution rates (number of fixations per year per site) of protein-coding genes and pseudogenes in the four chicken breeds with respect to orthologous sequences in RJF (GRCg6a)

|  | Daweishan | Hu | Piao | Wuding |
| --- | --- | --- | --- | --- |
| **Protein-coding gene** | 1.16e-09 | 1.41e-09 | 1.10e-09 | 1.10e-09 |
| **Pseudogene** | 1.04e-08 | 1.12e-08 | 1.02e-08 | 1.08e-08 |

### The GRCg6a assembly might be of an individual of *G. g. bankiva* origin

Although the RJF *G. g. spadiceus* subspecies is believed to be the major ancestor of domestic chickens all over the world [28], no high-quality genome of a *G. g. spadiceus* individual has yet been available. Thus, we would compare the gene compositions in our four indigenous chicken genomes with that of the RJF genome (GRCg6a). To infer the subspecies origin of the RJF individual and the indigenous chickens belonging to, we performed a principal component analysis (PCA) on the SNPs profiles of the RJF individual and populations of the five RJF subspecies (*G. g. Spadiceus, G. g. murphy*, *G. g. jabouillei*, *G. g. gallus* and *G. g. bankiva*) as well as of the four indigenous chickens (Methods and Materials). As expected, individuals of the four indigenous chicken breeds form a compact cluster with those of the *G. g. spadiceus* subspecies (Figure 4a), indicating that the four indigenous chicken breeds are indeed derived from *G. g. spadiceus*. Consistent with a previous report [28], individuals of *G. g. murphy* form a widely spread cluster that cannot be separated from the compact cluster formed by individuals of *G. g. jabouillei* (Figure 4a), suggesting the diversity of the individuals of *G. g. murphy* and possible admixture with *G. g. jabouillei.* Individuals of *G. g. gallus* form a cluster between the one formed by individuals of *G. g. jabouillei* and the one formed by individuals of *G. g. bankiva.* In agreement with the previous report [28], individuals of *G. g. bankiva* form a cluster that is farthest away from those formed by other subspecies and the indigenous chickens (Figure 4a), indicating that *G. g. bankiva* diverged earliest from the other subspecies. Interestingly, the sequenced RJF (GRCg6a) is separated far away from the cluster formed by the indigenous chickens and *G. g. spadiceus* individuals, and is closest to the cluster formed by *G. g. bankiva* individuals (Figure 4a), suggesting that it might be of *G. g. bankiva* origin. We also analyzed the genetic structures of the chickens (Methods and Materials). In agreement with the PCA result, GRCg6a has highly similar genetic structure to the individuals of *G. g bankiva* (Figures 4b-4d). Both *G. g. murphy* and *G. g. spadiceous* have diverse genetic structures (Figures 4b-4d) due to their broader geographic origins as previously indicted [28]. Hu and Piao chickens have mosaic genetic structures, while Daweishan and Wuding chickens have quite uniform genetic structures (Figures 4b-4d). The four indigenous chicken breeds have large genetic admixture from *G. g. spadiceous* but little from the other subspecies (Figures 4b-4d). These results further strengthen our conclusion that the four indigenous chicken breeds might be mainly derived from *G. g. spadiceous*, while the GRCg6a assembly belongs to an individual of *G. g. bankiva* origin. The latter conclusion might not be surprising given the fact that the sequenced RJF individual was from the UCD0001 line that was originated from a wild population from Malaysia [1], where *G. g. bankiva* inhabits [28].

**Figure 4.**
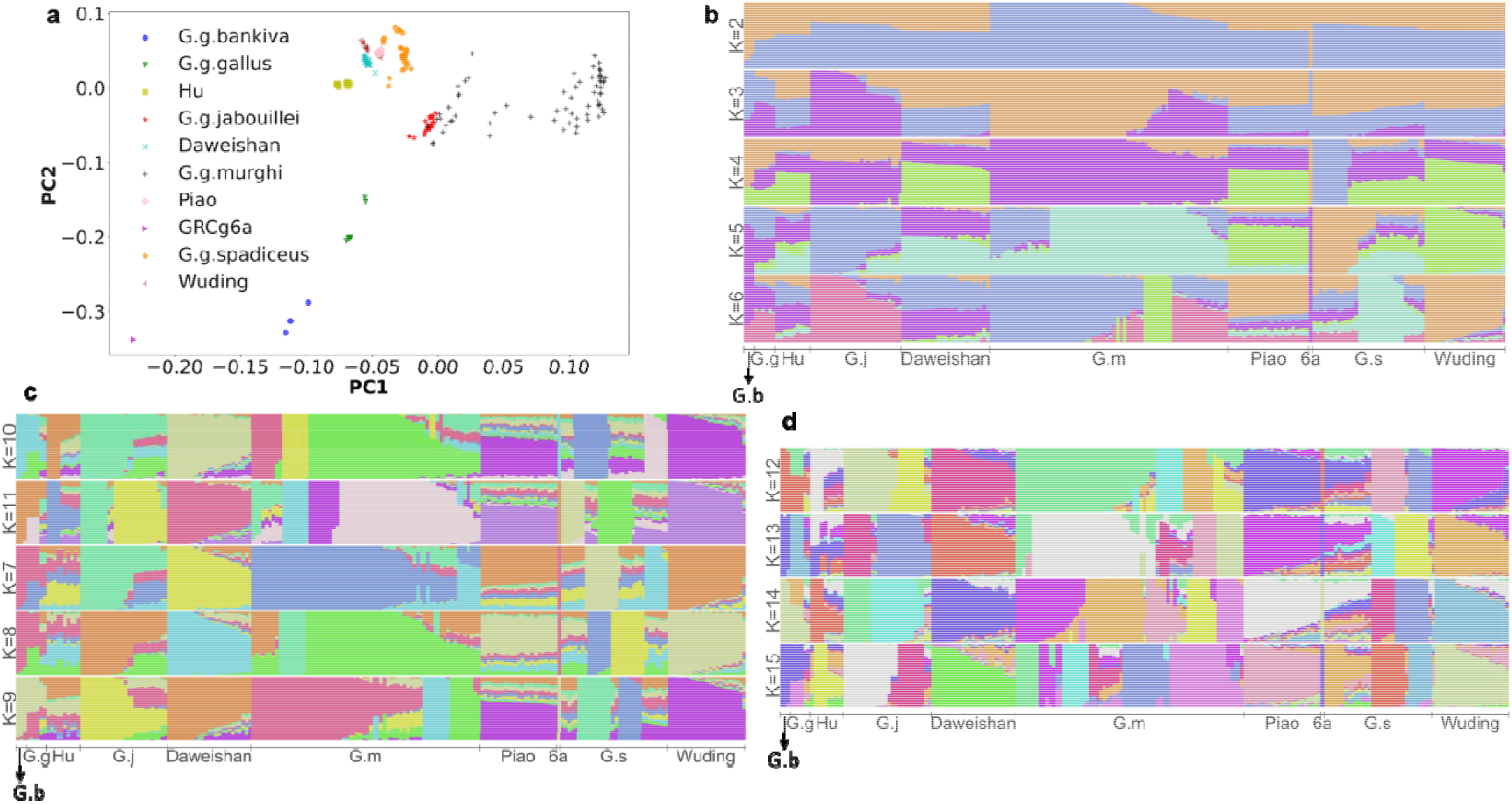
Analysis of frequency spectrums of SNPs. **a.** Principal component analysis of the RJF subspecies, our indigenous chickens and the RJF individual (GRCg6a) based on their SNP profiles**. b-d.** Genetic structures of the RJF subspecies, our indigenous chickens and the RJF individual (GRCg6a) estimated using the ADMIXTURE program for K=2, 3, …, 15.

### Chickens and the RJF have lost thousands of genes since their divergence

Based on our aforementioned results, it is reasonable to assume that our indigenous chickens and the RJF share a MRCA A1 before subspeciation, and the indigenous chickens share a MRCA A2 of *G. g. spadiceus* after subspeciation. There are two possible scenarios that the indigenous chickens and the RJF could be derived from the two MRCAs. In one scenario, a MRCA possessed at least the union of genes in the derived chickens plus functional versions of the intersection of unprocessed pseudogenes of all the derived chickens, and the derived chickens selectively lost genes via two unnecessarily exclusive forms of loss-of-function mutations, i.e., complete gene loss and pseudogenization, during the evolution and domestication processes. In the case of the four indigenous chickens, as illustrated in Figure 5a, their MRCA A2 would possess 20,760 genes (20,631 genes (Figure 1a) + 129 unprocessed pseudogenes (Figure S1b)), and Daweishan, Hu, Piao and Wuding chickens would lose functions of 3,042, 3,263, 3,049 and 3,114 genes, respectively, through pseudogenization (684, 556, 627 and 619) and complete gene loss (2,358, 2,707, 2,422 and 2,495) during their evolution and domestication processes. In the case of the four indigenous chickens and the RJF, their MRCA A1 would possess 21,972 genes (21,947 genes (Figure 1b) + 25 unprocessed pseudogenes (Figure S1a)), and the RJF and MRCA A2 would lose functions of 3,509 and 1,212 genes, respectively, through pseudogenization (462 and 0) and complete gene loss (3,047 and 1,212) during the subspeciation and evolution processes (Figure 5a). Moreover, from MRCA A1, Daweishan, Hu, Piao and Wuding chicken would lose function of additional 1,212 genes (Figure 5a). This explanation is in agreement with the earlier finding that chicken genome has undergone a large number of segmental deletions, resulting in a substantial reduction of genome sizes and the number of genes [1]. In the other scenario, the MRCA would possesses at most the intersection of genes in the derived chickens, and the derived chickens selectively gain genes during the evolution and domestication processes. In the case of the four indigenous chickens and the RJF, their MRCA A1 would possess at most 13,979 genes, and RJF, Daweishan, Hu, Piao and Wuding chickens would have gained 4,484, 3,739, 3,618, 3,732 and 3,667 genes, respectively, since their divergence (Figure 1b). However, this scenario is unlikely since there is no evidence of a large-scale introgression in the RJF and the indigenous chickens (Figure 1b).

**Figure 5.**
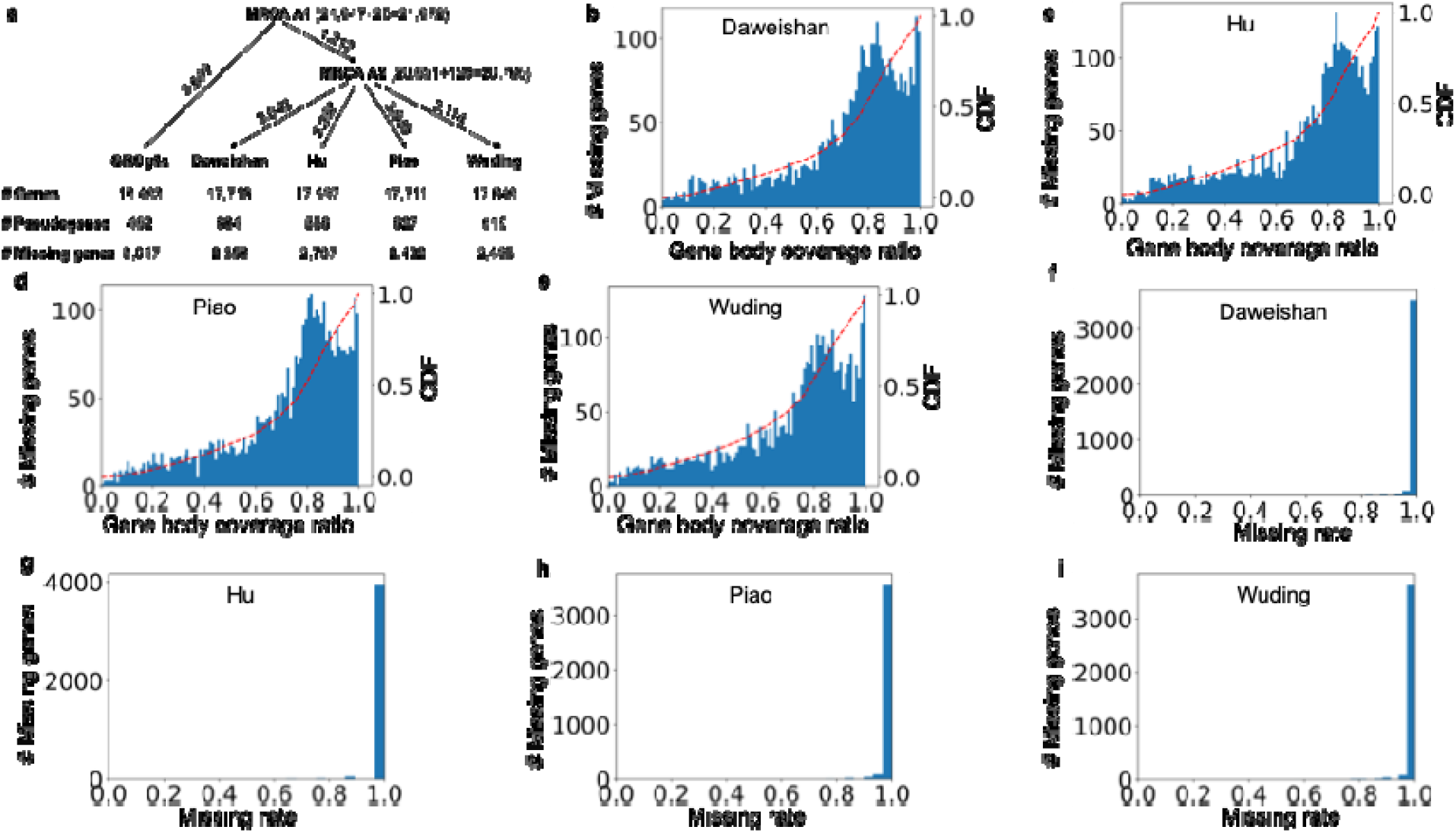
Evolutionary pattern of our four indigenous chickens and the RJF. **a.** A hypothetical scenario for loss-of-functions of the five chickens since their divergence from MRCA A1 and A2. **b-e.** Number of missing genes in each of our indigenous chicken genomes with the indicated reads coverage on their functional versions. The dashed red lines are the cumulative density function (CDFs) of coverage ratios. **f-i.** Number of missing genes with the indicated missing rate in the chicken populations.

Assuming MRCA A1 of the RJF and four indigenous chickens possessed 21,972 genes (Figure 5a), then, as the five chickens share 13,979 core genes (Figure 1b), the remaining 7,993 would lose functions through complete gene loss or pseudogenization in at least one of the five chickens (Table S27). Specifically, of the 7,993 dispensable genes, 1,583 (19.8%) are pseudogenized in at least one of the five chickens, while the remaining 6,410 (80.2%) are not pseudogenized in any of the five chickens, but are completely lost in at least one of the five chickens. Similar results are seen in the HiFi-reads assembled genomes. Specifically, with the similar assumptions, we found that MRCA A1 of the RJF and Huxu, Yueyang, Silkie and Hailanhe chickens had 23,628 genes, and 13,349 of which are core genes shared by the five genomes, implying that 10,279 genes in their MRCA A1 are dispensable genes (Table S28). Of the 10,279 dispensable genes, 1,401 (13.6%) are pseudogenized in at least one of the five chickens, while the remaining 8,878 (86.4%) are not pseudogenized in any of the five chickens, but are completely lost in at least one of the five chickens. For the convenience of discussion, we refer these completely lost genes as missing genes.

### Most missing genes have residual sequences in the indigenous chicken genomes but lose gene features in respective populations

To see whether the missing genes in the assembled genomes also are completely lost in their respective populations, we mapped short DNA reads from the individual chickens of a breed population to the functional versions of the missing genes in the corresponding assembly (Materials and methods). As shown in Figures 5b-5e, in each of our four indigenous chicken genomes, vast majority (99.9%) of the missing genes still have residual sequences in the individual genomes, covering up to 90% of their functional versions, while only few missing genes lack detectable residual sequences in the populations. These results strongly suggest that most of the missing genes were once functional in the ancestors, but lost gene features beyond recognitions, probably after becoming pseudogenes. In other words, their null-function mutations might have occurred earlier than most of the identified pseudogenes. Moreover, most (99%) of the missing genes in assembled indigenous chicken genome have a missing frequency > 99% in their respective populations, and only few are detected in all the re-sequenced individuals in respective populations (Figures 5f-5i). The high missing rate of the missing genes in the relevant populations suggests that loss-of-function (null) mutations are fixed in the populations, and thus might be under positive selection or artificial selection for their loss. Two examples of fixed loss-of-function mutations of missing genes in indigenous chicken populations are shown in Figure S3. In both cases, the residual sequences cover different parts of the functional versions of the missing genes in different breeds, with a missing match rate > 11% and all containing gaps (Figure S3). The lack of population data in most of the four HiFi-reads assemblies prevented us from conducting a similar analysis.

### Pseudogenes and functional versions of missing genes are preferentially located on micro-chromosomes and have high G/C contents

We analyzed the distributions of the 1,583 pseudogenes and the 6,410 missing genes in the chromosomes in each of our indigenous chicken genome. For missing genes in a chicken genome, we used its functional copy in another chicken for the analysis. Both the pseudogenes and functional versions of the missing genes are located in almost all the chromosomes in each of the four indigenous chickens (Figures 6a, 6b). However, the micro-chromosomes (chr14-chr39) and unplaced contigs have higher densities of both the pseudogenes and functional versions of the missing genes, harboring more than a third of the pseudogenes (39.4%-41.3%, Figure 6c) and more than half of functional versions of the missing genes (51.5%-54.6%, Figure 6d) though comprising only 13.5%-14.4% of the genomes. Likewise, both the ratio of the number of pseudogenes over the number of genes (Figure 6e) and the ratio of the number of functional versions of missing genes over the number of genes (Figure 6f) tend to be higher on the micro-chromosomes and unplaced contigs. Both the pseudogenes and functional versions of the missing genes (in other chickens) have a significantly higher G/C contents than true genes in each of the four indigenous chicken genomes (Figure 6g). Interestingly, the pseudogenes exhibit significantly higher G/C contents than functional versions of the missing genes for all the four chicken genomes except for the Hu chicken genome (Figure 6g). The same results were seen when the analyses were done separately on macro-chromosomes (chr1-chr5 and chrZ) (Figure S4a), intermediate-chromosomes (chr6-chr13 and chrW) (Figure S4b) and micro-chromosomes (chr14-chr39) (Figure S4c), to account for their different G/C contents [30]. It is unclear to us why functional copies of the missing genes have elevated G/C contents compared to true genes in the chicken genome. However, the elevated G/C contents in the pseudogenes compared with those in true genes and functional versions of the missing genes might be due to G/C-biased gene conversion during miotic recombination and DNA repairing [46] after purifying selection on the pseudogenes were relaxed. Consistently, pseudogenes in each indigenous chicken genome tend to have higher difference rates with their parental genes than do true genes with their template genes (Figure S5), indicating that pseudogenes have higher substitution rates. Since the contigs from the four HiFi-reads assembled genomes have not been assigned to chromosomes, we did not conduct the similar analysis.

**Figure 6.**
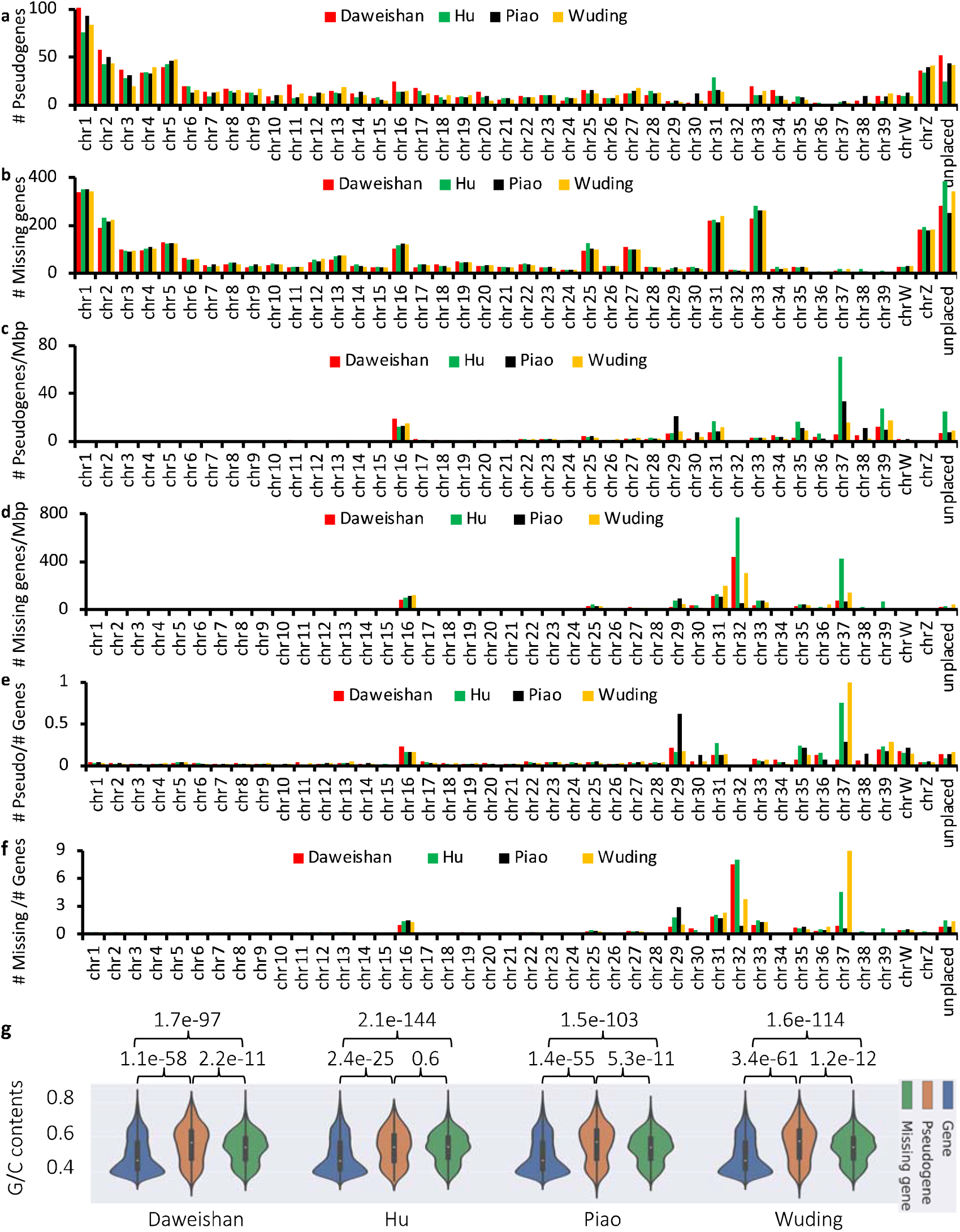
Distribution of pseudogenes and missing genes on each chromosome of the four indigenous chicken genomes. **a, b.** Number of pseudogenes (a) and missing genes (b) on each chromosome of the chicken genomes. **c, d.** Density of pseudogenes (c) and missing genes (d) on each chromosome of the chicken genomes. **e, f.** Ratio of the number of pseudogenes over the number of genes (e), and ratio of the number of missing genes over the number of genes (f), on each chromosome of the chicken genomes. **g.** Comparison of G/C contents of true genes, pseudogenes and missing genes in the chicken genomes. Statistical tests were done using one-tailed t-test.

### Loss-of-function mutations affect an array of important biological pathways of chickens

We analyzed the biological functions of the 7,993 dispensable genes in MRCA A1 that are either completely lost (6,410) or pseudogenized (1,583) in at least one of our four indigenous chickens and RJF (Figure 5a). To this end, we performed a two-way hierarchical clustering on the 7,993 dispensable genes and the five chickens based on the occurring patterns of these dispensable genes in the five chicken genomes based on Euclid distances using the UPGMA method. The dispensable genes form distinct clusters along the clustering hierarchy (Figure 7a). Based on the distinct features of clusters, we divided them into 31 exclusive clusters as described in Table S27. Although only 1,567 (19.6%) of the dispensable genes have GO term assignments to their functional parental genes in GRCg6a/GRCg7b/GRCg7w or humans, most clusters (27/31, 87.1%) containing genes related to important biological pathways (Figure 7a, Table S27).

**Figure 7.**
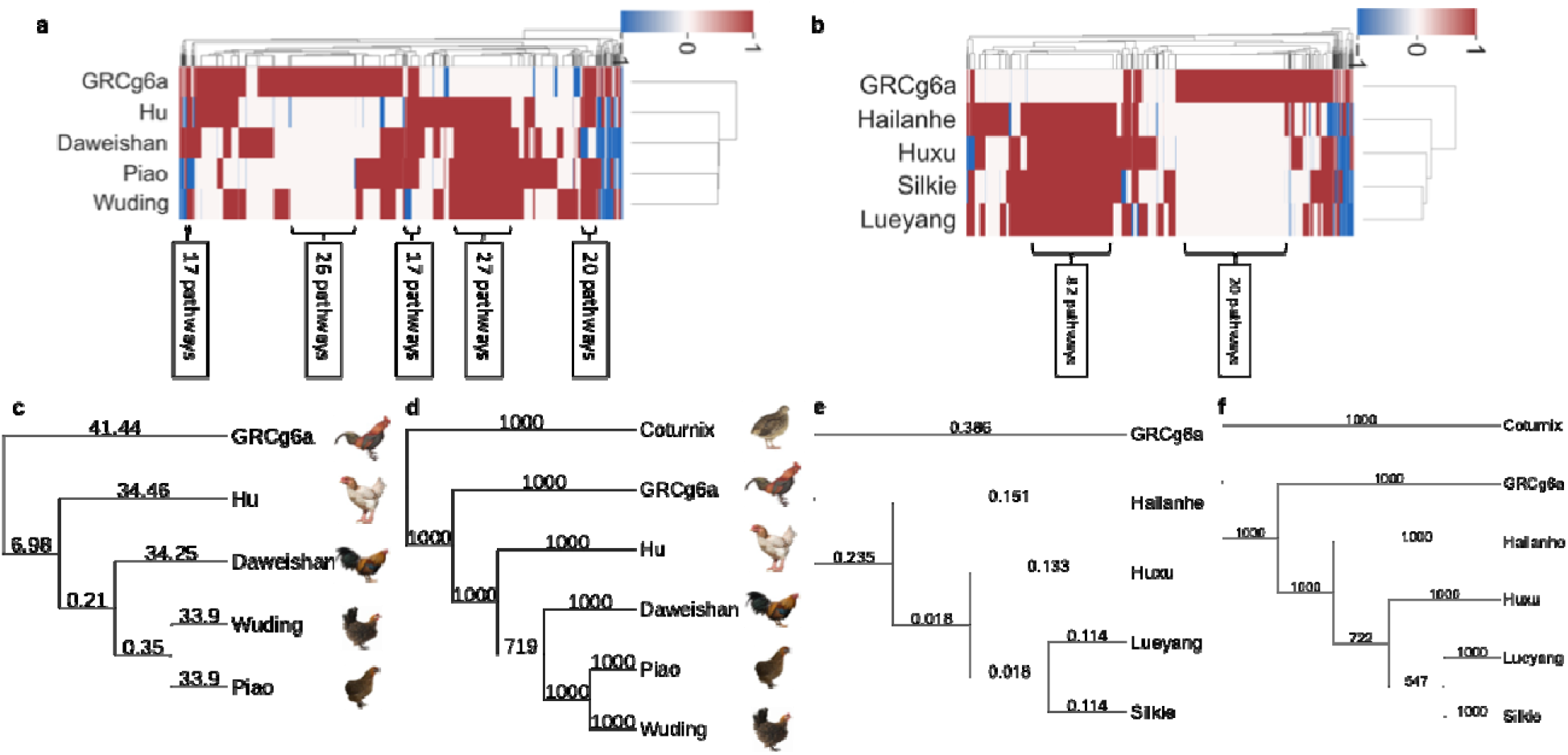
Evolutionary relationships of each chicken breed. **a.** Heatmap of two-way hierarchical clustering of the dispensable genes in MRCA A1 that are either completely lost or pseudogenized in at least one of our four indigenous chickens and RJF based on their appearance as an intact form (1, brown), absence (0, white) and as a pseudogenized form (-1, blue) in the five chicken genomes. **b.** Heatmap of two-way hierarchical clustering of the dispensable genes in MRCA A1 that are either completely lost or pseudogenized in at least one of the four HiFi-reads assembled genomes and RJF based on their appearance as an intact form (1, brown), absence (0, white) and as a pseudogenized form (-1, blue) in the five chicken genomes. **c.** Neighbor-joining phylogenetic tree of our four indigenous chickens and RJF, constructed using the occurring patterns of the dispensable genes in their genomes. The numbers on the branches are Euclid distance between the pattern vectors. **d.** Neighbor-joining phylogenetic tree of our four indigenous chickens and RJF, constructed using the 6,744 essential protein-coding genes in their genomes and the quail genome. The numbers on the nodes are bootstrapping value for 1,000 repeats. **e.** Neighbor-joining phylogenetic tree of the four HiFi-reads assembled genomes and RJF, constructed using the occurring patterns of the dispensable genes in their genomes. The numbers on the branches are Euclid distance between the pattern vectors. **f.** Neighbor-joining phylogenetic tree of the four HiFi-reads assembled genomes and RJF, constructed using the 6,323 essential protein-coding genes in their genomes and the quail genome. The numbers on the nodes are bootstrapping value for 1,000 repeats.

For instance, cluster 29 containing 241 genes that are completely lost or pseudogenized in Daweishan chicken but functional in other four genomes are involved in 20 pathways, including Thiamin metabolism pathway which is enriched (p-value = 1.03e-02, Fisher’s Exact test). Thiamin is a key cofactor in carbohydrate metabolism and ATP production. Disruption of this pathway may reduce metabolic efficiency and energy generation, potentially constraining growth. This may contribute to the small body size observed in Daweishan chicken; Cluster 28 containing 285 genes that are completely lost or pseudogenized in Hu chicken but functional in other four chickens are involved in 25 pathways, including Phenylethylamine degradation pathway which is enriched (p-value = 2.72e-02). Phenylethylamine is involved in neuromodulation and metabolic regulation. Altered degradation may influence feeding behavior and energy balance, potentially contributing to the larger body size of Hu chicken; Cluster 27 containing 221 genes that are completely lost or pseudogenized in Piao chicken but functional in other four chickens are involved in 17 pathways, including Glutamine glutamate conversion pathway (p-value = 2.40e-02) and Mannose metabolism pathway (p-value = 2.87e-02) which are enriched. These pathways are important for cell proliferation and protein glycosylation. Their disruption may affect signaling and cellular dynamics during embryogenesis, potentially impairing posterior axis development and contributing to the rumpless phenotype of Piao chicken; Cluster 26 containing 217 genes that are completely lost or pseudogenized in Wuding chicken but are functional in other four chickens are involved in 17 pathways related to neuromuscular signaling, cell motility and muscle development (e.g., Rho GTPase, G-protein and growth factor signaling). Such changes may reshape muscle function and coordination, potentially contributing to enhanced running ability of Wuding chicken.

Cluster 30 containing 1,071 genes that are functional in all the four indigenous chicken genes but are completely lost or pseudogenized in the RJF are involved in 27 pathways, including ATP synthesis pathway which is enriched (p-value = 2.29e-02). As ATP production is central to cellular energy metabolism, these differences may reflect distinct energy utilization strategies between wild and domesticated chickens, potentially associated with artificial selection for growth and productivity; Cluster 1 containing 1,303 genes that are functional in the RJF but completely lost or pseudogenized in all the four indigenous chickens are involved in 26 pathways related to development, immunity, stress response, and behavior. These pathways are essential for environmental adaptation and physiological regulation. Their loss in domesticated chickens suggests a broad reorganization of biological systems during domestication, likely reflecting reduced environmental pressures and selection for production-related traits.

Although further functional validation is required, the importance of these affected pathways suggests that loss-of-function mutations might have shaped the traits of the indigenous chickens for them to adapt to their ecological and domesticated conditions and for the RJF to adapt to its ecological niche. Similar results were obtained for the 10,279 dispensable genes in MRCA A1 of RJF and Huxu, Yueyang, Silkie and Hailanhe chickens (Figure 7b, Table S29).

### Occurring patterns of loss-of-function mutations reflect evolutionary history of the chickens

The patterns of loss-of-function mutations of genes in the chicken genomes might provide hints to mutation orders during chicken evolution and domestication. Most pseudogenes in the GRCg6a genome are completely lost in our four indigenous chicken genomes (Figure 7a), suggesting that the indigenous chickens completely lost these genes after their divergence from the MRCA A1, during their domestication process and/or subspeciation of *G.g. spadiceous*, while GRCg6a is still in the process of completely losing these genes. In contrast, most pseudogenes in the four indigenous chicken genomes have intact copies in GRCg6a (Figure 7a), suggesting that these GRCg6a genes might lose their functions in the indigenous chickens after their separation from MRCA A1, during the subspeciation and domestication processes, and might be in the process of being completely lost.

To see whether the occurring patterns of the complete gene loss and pseudogenization in the five chicken genomes reflect their evolutionary relationships, we constructed a neighbor-joining (NJ) phylogenetic tree using the occurring patterns in the five chicken genomes of the 7,993 MRCA A1 genes (Table S27) that are either completely lost or pseudogenized in at least one of the five genomes. As shown in Figure 7c, consistent with the UPGMA tree rooted with the RJF (Figure 7a), Wuding and Piao chickens form a clade that is joined by Daweishan chicken, and the resulting cluster is joined by Hu chicken. The tree is also consistent with the NJ tree constructed using 6,744 essential protein-coding genes in the five chicken genomes and quail (*Coturnix jcponica*) with quail as the root (Figure 7d, Table S30). Therefore, the occurring patterns of complete gene loss and pseudogenization in the five chickens segregate them in the way by their evolutionary relationships, and thus, reflect their evolutionary relationships.

Similar results were obtained for the HiFi-reads assembled genomes. Specifically, the NJ tree rooted with the RJF constructed on the base of occurring patterns of the 10,279 dispensable genes in MRCA A1 of the RJF and Huxu, Yueyang, Silkie and Hailanhe chickens (Table S28), is consistent with the UPGMA tree rooted with the RJF (Figure 7b) in that Lueyang and Silkie chickens form a clade that is joined by Huxu chicken, and the resulting clade is joined by Hailanhe chicken. The tree is also consistent with the NJ tree constructed using 6,323 essential protein-coding genes in the five chicken genomes and quail (*Coturnix jcponica*) (Table S31) with quail as the root (Figure 7f), suggesting again that pseudogenization in each chicken genome could reflect their evolutionary relationships. This result is in contrast to the earlier reports that loss-of-function mutations failed to segregate between wild and domestic chickens, and thus selection for loss-of-function mutations had a little role in chicken domestication and evolution [12, 13].

## Discussion

We find large numbers of pseudogenes in each chicken genomes analyzed, regardless of the methods used for their assembling. Most of the pseudogenes in each chicken genome are unprocessed and unitary, while only a small number of them are processed. This finding is consistent with the previous results [47, 48], presumably because the chicken’s LINE1 like CR1 (L1) elements lack retro transposase activity [47, 48]. However, the large number of unprocessed pseudogenes that we found in each chicken genome is in stark contrast to the findings in some mammalian species such as humans and mice. For example, a previous study found that only a few dozen unprocessed pseudogenes were found in human population [49, 50], not to mentioning in a single human individual genome. In a more recent study [51], 165 and 303 unprocessed pseudogenes were found in large mouse and human populations, respectively. However, these numbers are still much smaller than those (3,069, pseudogenes in at least one of the genomes) that we found in only nine chicken genomes. Thus, we observed a larger scale of unprocessed pseudogenization in chickens than in mice and humans.

Our results strongly suggest that most of the pseudogenes lose their protein-coding functions. First, the pseudogenes tend to have elevated G/C contents and higher dN/dS ratios compared with true genes, no matter where the first pseudogenizations occur along the CDSs of their parental genes, indicating that they are no longer under purifying selection but might be under positive selection for loss of function. Second, in true genes, synonymous mutations are largely uniformly distributed along the CDSs, but their occurrences decrease at the two-ends of the CDSs, suggesting that both ends are under stronger purifying selection. However, such stronger purifying selection is relaxed on the two ends of pseudogenes. Third, although most pseudogenes have transcripts, very few have isoforms that skip the exon with the first pseudogenization mutations, and thus, unlikely producing functional peptides. Finally, there are two scenarios for a pseudogene to arises: 1) when the function of the gene is no longer needed; and 2) after a gene duplication event, removal of a redundant copy is beneficial. We find that most of the pseudogenes in the chickens are unitary, and thus, are not related to gene duplications. Therefore, functions of most parental genes are lost in the genomes that harbor the pseudogenes.

Moreover, we find that the compositions of protein-coding genes in the chicken genomes are highly variable even though they harbor similar number of genes. For example, although the Daweishan and Piao chicken genomes encode almost the same number of genes (17,718 vs 17,711), they share only 91% of their genes even though they have diverged for only a few thousand years [28]. These results are in stark contrast with the observation that even some closely-related different species such as humans and chimpanzees, which share almost the same sets (98%) of genes, though being split at least 6-7 million years ago [36]. Both the unexpectedly large number of pseudogenes and the highly variable compositions of the proteomes in the chicken genomes strongly suggest that chickens have undergone dramatic changes in their proteomes in the past 500,000 years of evolution and domestication.

We confirm that the four indigenous chickens from Yunnan are mainly derived from the *G. g. spadiceous* subspecies, and infer for the first time that the sequenced RJF (GRCg6a) is of *G. g. bankiva* origin, providing an ideal case to investigate the evolution of gene composition in chicken genomes. Specifically, it has been reported that these indigenous chickens and the RJF might share their MRCA A1 50,000-500,000 years ago [28]. There are two possible scenarios that the highly variable proteomes in the five chicken genomes could arise: 1) their MRCA A1 only possessed the intersection of their genes, and each derived chicken selectively gained thousands of new genes; and 2) their MRCA A1 harbored the union of their genes plus functional versions of the intersection of their unprocessed unitary pseudogenes genes, and each derived chicken selectively lost thousands of genes. Our results are against scenario 1 but favor scenario 2. First, the RJF and the four indigenous chickens have little gene introgression from other RJF subspecies (Figures 4b-4d). Thus, it is unlikely that the five chickens have gain large numbers of genes from other subspecies. Second, most of the unprocessed unitary pseudogenes in each chicken genome arose quite recently during the processes of domestication (∼10,000 years ago) and evolution during (50,000-500,000 years ago) and after subspeciation (10,000-50,000 years ago) (Figures 3m and 3n), and are still in the process of being completely lost, and hence, they might be once functional in recent ancestors. Third, although all the missing genes in a genome have lost all gene features, most of them have residual sequences left in the genomes, strongly suggesting that they were also once functional in recent ancestors. The missing genes might have become pseudogenes earlier than most of the predicted pseudogenes, because the former have lost all gene features beyond being identified as even pseudogenes, while the latter still possess few to be recognized as pseudogenes. In addition, it has been shown that the RJF genome has undergone a large-scale of segmental deletions, resulting in a substantial reduction of the number of genes [1].

Although 7,993 of the 21,947 genes estimated in the MRCA A1 of the RJF and our four indigenous chicken genomes are dispensable, many of them are involved in important biological processes. Thus, their selective retainment or loss in a genome might be beneficial to the chicken in its unique natural or domestic conditions. More specifically, although the thousands of genes in the RJF genome that are either completely lost or pseudogenized in the four indigenous chicken genomes might be essential for RJF to live in its unique ecological niche, loss-of-function mutations of these genes in the indigenous chickens might be beneficial for them to live in domestic conditions. Similarly, although the thousands of genes in our four indigenous chicken genomes that are either completely lost or pseudogenized in the RJF genome might be essential for the indigenous chickens to live in their domestic conditions, loss-of-function mutations of these genes in the RJF might be beneficial for it to live in its unique ecological niche. These conclusions are supported by the results drawn from joint analysis of the four HiFi-reads assembled chicken genomes and the RJF genome.

Our results strongly suggest that loss-of-function mutations via complete gene loss and pseudogenization are a result of natural and artificial selection. First, unlike synonymous mutations along the true genes and pseudogenes, which are largely uniformly distributed along the CDSs as expected for neutral mutations, the most upstream pseudogenization mutation in pseudogenes is strongly biased to the two ends of parental CDSs, particularly, the 5’-ends. Such biases would facilitate eliminating the functions of parental genes. It is well-known that a promoter can extend into the 5’-end of the CDSs, thus mutations in the region may disrupt the promoter [52]. Moreover, the closer a pseudogenization mutation is toward the 5’-end of the CDSs, the greater impact of the mutation could have on the gene function and the more likely the gene would lose its function. Although pseudogenizations at the 3’-ends of CDSs can potentially produce at least partially functional proteins, this is unlikely for at least most of the pseudogenes that we found in the chicken genomes. This is because dN/dS ratios for pseudogenes with the first pseudogenization sites occurring in the last 10% and in the first 10% lengths of the CDSs are not significantly different, and both are significantly higher than those for true genes. In other words, pseudogenization mutations in the 3’-ends of CDSs can be as effective as those in the other parts of the CDSs to eliminate the functions of genes. We found that 3’-ends of CDSs might harbor miRNA binding sites, and pseudogenization mutations could disrupt such binding sites, which thereby might change post-transcriptional regulation, and thus, the functions of genes.

Second, most pseudogenization mutations are fixed in our indigenous chicken populations (Figures 3a-3d), and thus they are likely under strong positive selection or artificial selection for null alleles. Third, pseudogenes mainly arose in the past 500,000 years (Figures 3m and 3n), correspond to the domestication period (past 10,000 years) [29], subsequent evolutionary period (10,000-50,000 years ago) after subspeciation, and subspeciation period (50,000-500,000 years ago) [28]; and thus, they are mainly resulted from recent natural selection and artificial selection. Unlike completely lost genes, pseudogenes have not had enough time to be fully degraded after they are no longer subject to negative selection. Of course, with time the pseudogenes without any other functions will be eventually degraded beyond recognition and become missing. Fourth, most of the missing genes in each of our assembled indigenous chicken genome also are missing in the corresponding population (Figures 5f-5i), i.e., the null alleles are fixed, suggesting that the missing genes might be under strong positive or artificial selection for losing. Finally, the occurring patterns of the dispensable genes in the MRCA A1 segregate chickens and the RJF in the exact same way as the phylogenetic tree of the chickens constructed using essential avian protein-coding genes (Figures 7c-7f). Taken together, these results strongly suggest that loss-of-function mutations via pseudogenization and complete loss of thousands of genes in RJF and the chickens since their divergence might play critical roles in chicken evolution and domestication. This conclusion is in contrast to an earlier report that loss-of-function mutations play a little role in chicken domestication [12]. Complete gene loss and pseudogenization are not unnecessarily exclusive forms of loss-of-function mutations. Once a gene is pseudogenized, it will be rapidly degraded as purifying selection on the pseudogene is relaxed (Figures 2g and 2h).

It is worth pointing out that although it has been shown that deleterious mutations might play roles in the domestication and evolution of plants [53, 54] and animals [55, 56], lost-of-function mutations are not necessarily deleterious. In fact, it has been well documented that loss of certain genes might be the results of adaptation of birds for flight [21, 57–59], of beef cattle for meat production [25], and of humans for new abilities [49]. It has been proposed that loss-of-function mutations may be an important factor in rapid evolution as occurred during domestication—the “less is more” hypothesis [24], which has since gained substantial evidence supports [14, 15, 23, 60–67]. Consistently, in the present study we found that dozens (6%-7%) of pseudogenes arose in each of the four indigenous chicken breeds during the past 10,000 years of domestication process. Thus, the earlier conclusion that fixation of null alleles is not a common mechanism for phenotypic evolution in chicken domestication [12, 13] might be incorrectly drawn because of the low quality of then available chicken genome assemblies, leading to the failure to detect inactivating mutations such as large scale pseudogenizations and high variation of gene presence and absence [68].

## Materials and Methods

### Chicken population

The GRCg6a, and quail (*coturnix japonica*) genomes and annotation files were downloaded from the NCBI Genbank with accession numbers GCF_000002315.6, and GCF_001577835.2, respectively. Our previously assembled four indigenous chicken genomes were downloaded from the NCBI Genbank with the BioProject number PRJNA865263. All the Illumina short DNA sequencing reads and RNA-seq reads of different tissues of the four indigenous chickens were downloaded from the NCBI SRA database with accession number PRJNA865247. HiFi-reads assembled genomes of four chicken breeds were downloaded from NCBI Genbank with accession numbers GCA_024206055.2 (Huxu chicken), GCA_034509845.1 (Hailanhe chicken), GCA_034509865.1 (Silkie chicken) and GCA_034509885.1 (Lueyang chicken).

We downloaded the re-sequencing data of different RJF subspecies from the ChickenSD database (http://bigd.big.ac.cn/chickensd/) with accession numbers listed in [28]. All the re-sequencing data of the indigenous chickens were downloaded from the NCBI SRA database with the accession number PRJNA893352. The sequences of essential avian proteins were obtained from the BUSCO aves_odb10 database [37].

### Protein-coding gene and pseudogene annotation

We used a combination of homology-based and RNA-seq-based method to annotate the protein coding genes and pseudogenes as previously described [31]. Briefly, for the homology-based method, we firstly collected all protein-coding genes and pseudogenes in the GRCg6a, GRCg7b and GRCg7w genome as the templates, and then mapped all the CDS isoforms or exons of these genes and pseudogenes to each of the assembled indigenous chicken genomes using Splign (2.0.0) with default settings [69]. For each template gene whose CDSs can be mapped to an assembled genome, we concatenate all the mapped CDSs to form a putative full-length CDS. We predict corresponding sequence to be an intact gene if and only if the length of the putative full-length CDS is an integer time of three and the last three nucleotides form a stop codon but there are no stop codons in the other part of the putative full-length CDS. If the CDSs of a template gene can be mapped to multiple loci in an assembled genome, we consider the locus with the highest mapping identity. If the putative full-length CDS contains a premature stop codon (nonsense mutation), or its length is not an integer time of three (ORF shift mutation), we verify the mutations by mapping the short DNA reads from the same individual to the genomic locus using bowtie (2.4.1) [70] with no gaps and mismatches permitted. If the locus can be completely covered by at least 10 short reads at each nucleotide position, we consider the loss-of-function mutation (nonsense mutation or ORF shift mutation) is fully supported by the short reads, and predict the corresponding sequence to be a pseudogene. Otherwise, we consider the loss-of-function mutation is not supported by the short reads, and call the sequence a partially supported gene, because the loss-of-function mutation might be artificially caused by errors of the long reads that could not be corrected by assembly pipeline.

### Single nucleotide variants calling

We mapped short DNA reads from each individual chicken to the GRCg7b reference genome using Bowtie (2.4.1), and called SNVs and small indels in each individual chicken using GATK (4.1.6) [71].

### Calculation of allele frequencies of pseudogenes

We computed allele frequencies of the first pseudogenization mutation of each pseudogene in each chicken breeds using GATK (4.1.6) [71] based on call SNVs and indels.

### Neighbor-joining tree construction

We mapped the essential avian proteins to each of the five chicken’s CDSs as well as the quail’s CDSs using blastx (2.11.0) [72]. We selected the genes with greater than 70% sequence identity with the essential avian proteins in each of the genomes to construct a neighbor-joining tree. Since it is hard to make multiple alignments for very long sequences, we evenly divided the genes in each bird into multiple groups (each contains about 100 genes). We then aligned sequences of the same group in the genomes using Clustal Omega (1.2.4) [73]. We finally concatenated the many multiple alignments with a fixed order and constructed a consensus neighbor-joining trees with 1,000 rounds of bootstrapping using Phylip (3.697) [74].

### PCA and population structure analysis

We used the SNPs called in each individual chicken of each population to perform the PCA and population genetic structure analysis. PCA was performed using PLINK (1.90) [75] with the default settings, and population genetic structure analysis was inferred using ADMIXTURE (1.3.0) [76] with K=2, 3, …, 15 using the default settings.

### Prediction of miRNA binding sites

For each pair of pseudogene and its parental gene, we scanned their CDSs and 1,000 bp downstream sequences as putative 3’-UTRs for miRNA binding sites using RNAhybrid (2.1.2) [77]. The miRNAs predicted in the genome harboring the pseudogene are used as the database for the scanning. We consider the putative binding sites with a p-value<0.05.

### Calculation of gene body coverage ratio and missing rates of missing genes

For each assembled indigenous chicken breed genome, we collected functional version (reference genes) of its missing genes from either the RJF genome (GRCg6a) or any of the other three indigenous chicken genomes. We mapped the re-sequencing short reads of each individual chicken of each breed (n= 25, 10, 23 and 23 for Daweishan, Hu, Piao and Wuding, respectively) to the reference genes for the breed using Bowtie (2.4.1) [70] allowing no mismatch and gaps. For each missing gene in the assembled genome of a breed, we computed the gene body coverage ratio as the average length of the reference gene body covered by reads among all the individuals of the breed over the length of the reference gene body. We also computed missing rate for each missing gene in the assembled genome of a breed as the ratio of the number of individuals whose re-sequencing reads cannot fully cover the reference gene body over the number of total individuals of the breed.

### Estimation of substitution rates and divergent times of protein-coding genes/pseudogenes

We estimated the substitution rates of a gene/pseudogene since its divergence from the last common ancestral gene shared with the RJF (GRCg6a) by using the Kimura’s two-parameter (K2P) model [44]. Specifically, we aligned the CDSs of protein-coding genes and exons of pseudogenes in each indigenous chicken genome to the orthologous CDSs/exons of the GRCg6a assembly using blastn [72] by default settings. We counted the number of transition mutation sites and transversion mutation sites in each gene/pseudogene based on the alignments. We mapped the short reads of from the individuals of each breed population to the alleles using Bowtie (2.4.1), and filtered out those with a frequency < 0.2 in the corresponding breed population. We calculated the transition rate (*P*) and transversion rate (*Q*) of a gene/pseudogene as:

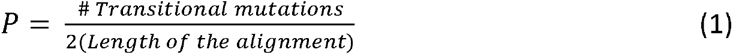

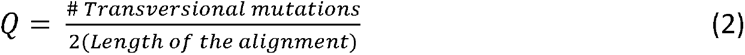

The factor 2 in the denominators in formulas (1) and (2) is to account for the fact that the diploid genomes are assembled as haploid mosaic ones. The number of substitutions per site per generation in the gene/pseudogene is computed as:

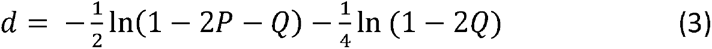

The mean of the number of substitutions of all genes/pseudogenes in a breed is defined as:

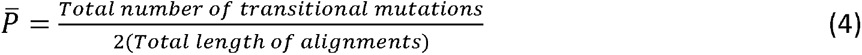

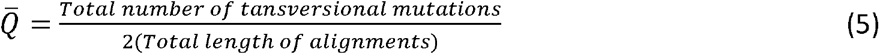

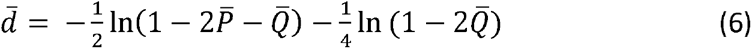

The substitution rate λ of genes/pseudogenes is estimated as:

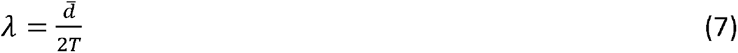

assuming a generation time of one year and the divergent time (*T*) between the indigenous chickens and the RJF (GRCg6a) about 100,000 years, an intermediate value of the estimated subspecialties time (50,000-500,000 years ago) of the *G. gallus* subspecies [28]. The time of divergence *t* between a gene/pseudogene and the last common ancestral gene shared with GRCg6a is estimated as:

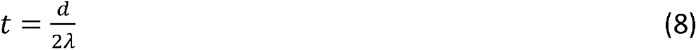

where *d* is the number of substitutions in the gene/pseudogene.

## Acknowledgments

This work was supported by the National Natural Science Foundation of China (U2002205 and U1702232), Yunling Scholar Training Program of Yunnan Province (2014NO48), Yunling Industry and Technology Leading Talent Training Program of Yunnan Province (YNWR-CYJS-2015-027), Natural Science Foundation of Yunnan Province (2019IC008 and 2016ZA008), and Department of Bioinformatics and Genomics of the University of North Carolina at Charlotte.

## Author contributions

JJ, CG and ZS supervised and conceived the project; KW, XG, TD, SY^2^, ZX, YL, ZJ, JZ, RZ, XZ, DG, LL, QL and DW collected tissue samples and conducted molecular biology experiments; SW and SY^1^ assembled and corrected the genomes; SW and ZS performed data analysis; and SW and ZS wrote the manuscript.

## Data availability

The annotation code and pipeline description are available at https://github.com/zhengchangsulab/HRannot

