## Supplementary Figures for "Large scale loss-of-function mutations during chicken evolution and domestication"

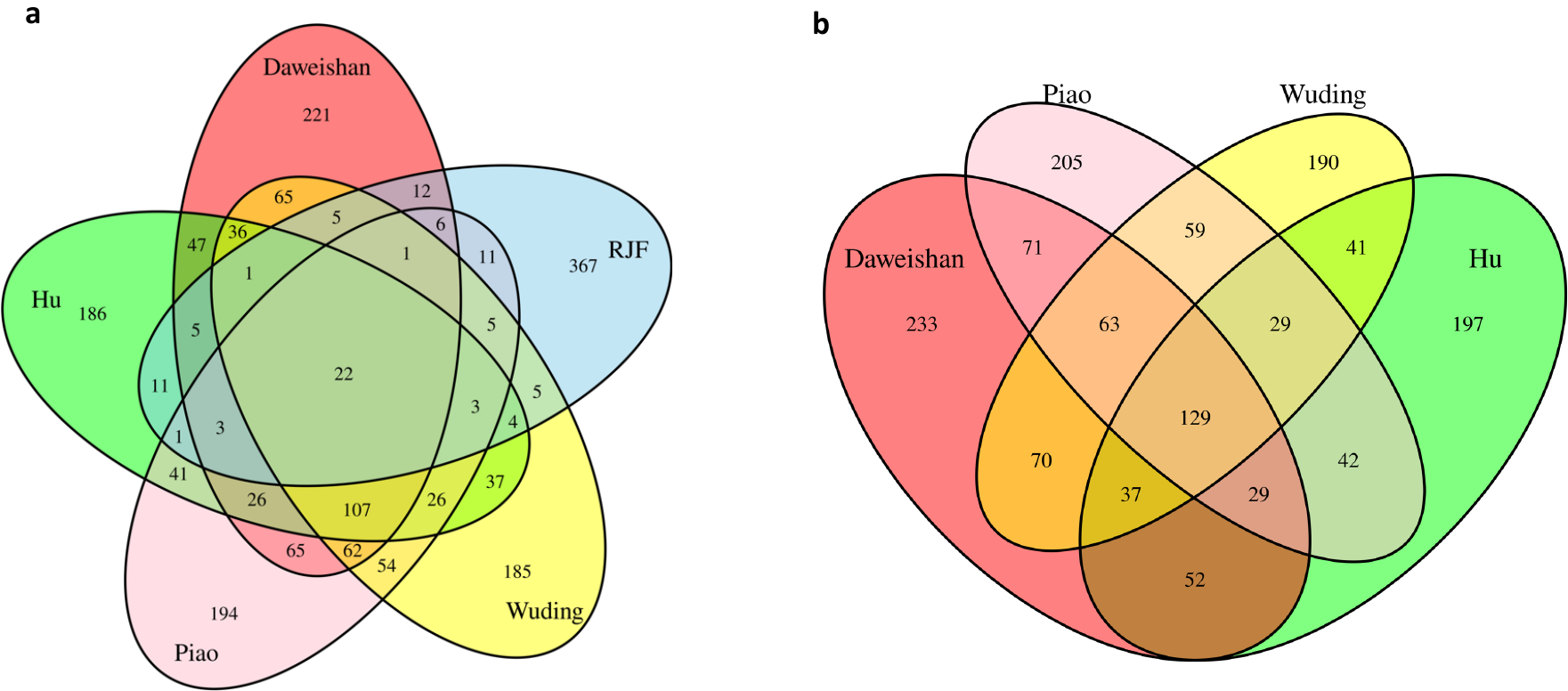

**Figure S1.** Venn diagrams of unprocessed pseudogenes among the five or four chickens. **a.** Venn diagram of unprocessed pseudogenes among the five chickens. **b.** Venn diagram of unprocessed pseudogenes among the four chickens.

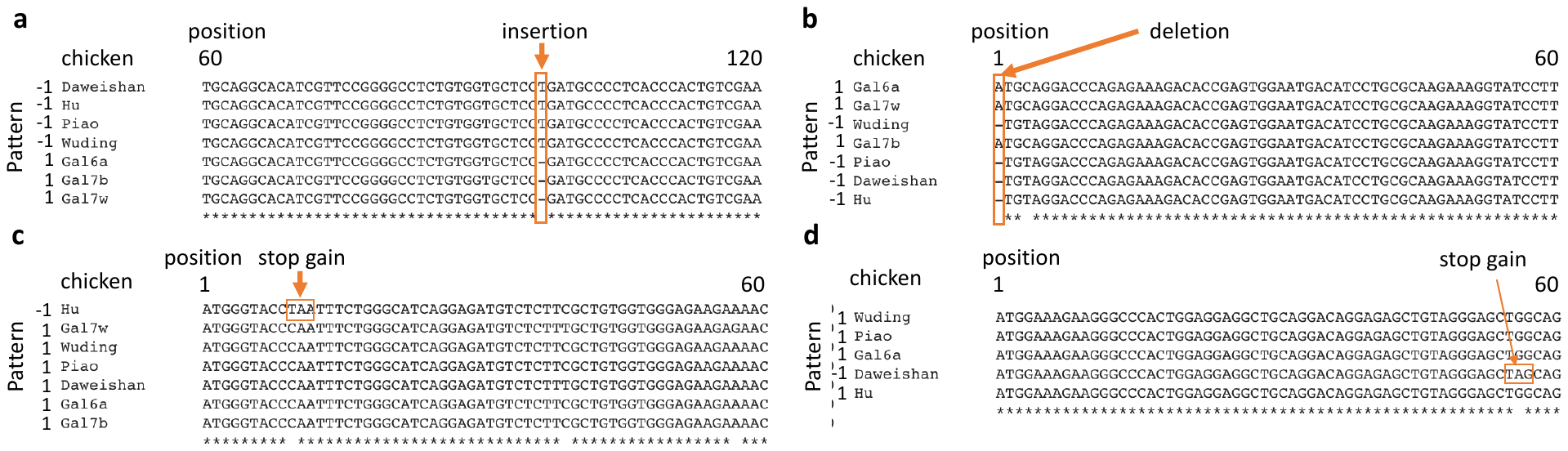

**Figure S2.** Examples of fixed or nearly fixed pseudogenes in chicken populations. **a.** The *OAZ*2 gene (CDS length=573bp), which is functional in GRCg6a, CRCg7b and CRCg7w, is pseudogenized and fixed in the four indigenous chickens, due to an insertion of a “T” after position 96 of the CDS, resulting an ORF shift. **b.** The *PDCL*3 gene (CDS length=723bp), which is functional in GRCg6a, CRCg7b and CRCg7w, is pseudogenized and fixed in the four indigenous chickens, due to deletion of an “A” at the first position of the CDS, resulting an ORF shift. **c.** The *PRDM*16 gene (CDS=3,828bp), which is functional in the six other chickens, is pseudogenized and fixed in the Hu chicken population, due to a “C to T” substitution at position 10 of the CDS, resulting a stop-gain. **d**. The *ZNF*408 gene (CDS=603bp), which is functional in the three other indigenous chickens and GRCg6a but missing in GRCg7b and GRCg7w, is pseudogenized and almost fixed (allele frequency=0.88) in the Daweishan chicken population, due to a “G to A” substitution at position 56 of the CDS, resulting a stop-gain.

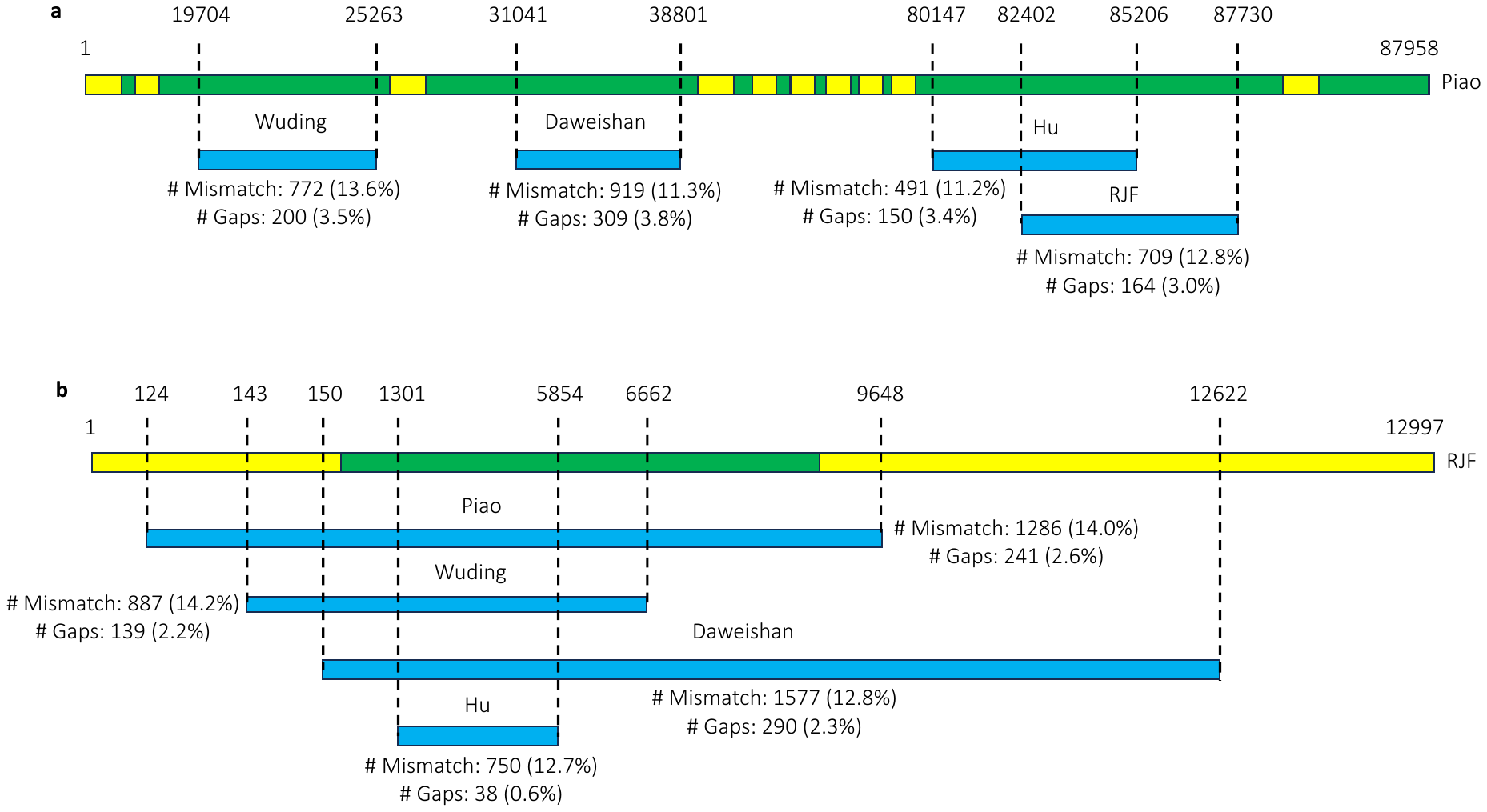

**Figure S3.** Examples of fixed missing genes in chicken populations. **a.** Gene SND1 on chr1 is present in Piao chicken populations, but missing in Daweishan, Hu, Wuding and RJF populations. **b.** Gene LOC107055343 on chr28 is present in RJF populations, but missing in Daweishan, Hu, Piao and Wuding populations. In the figures, the yellow parts represent the exons of the gene.

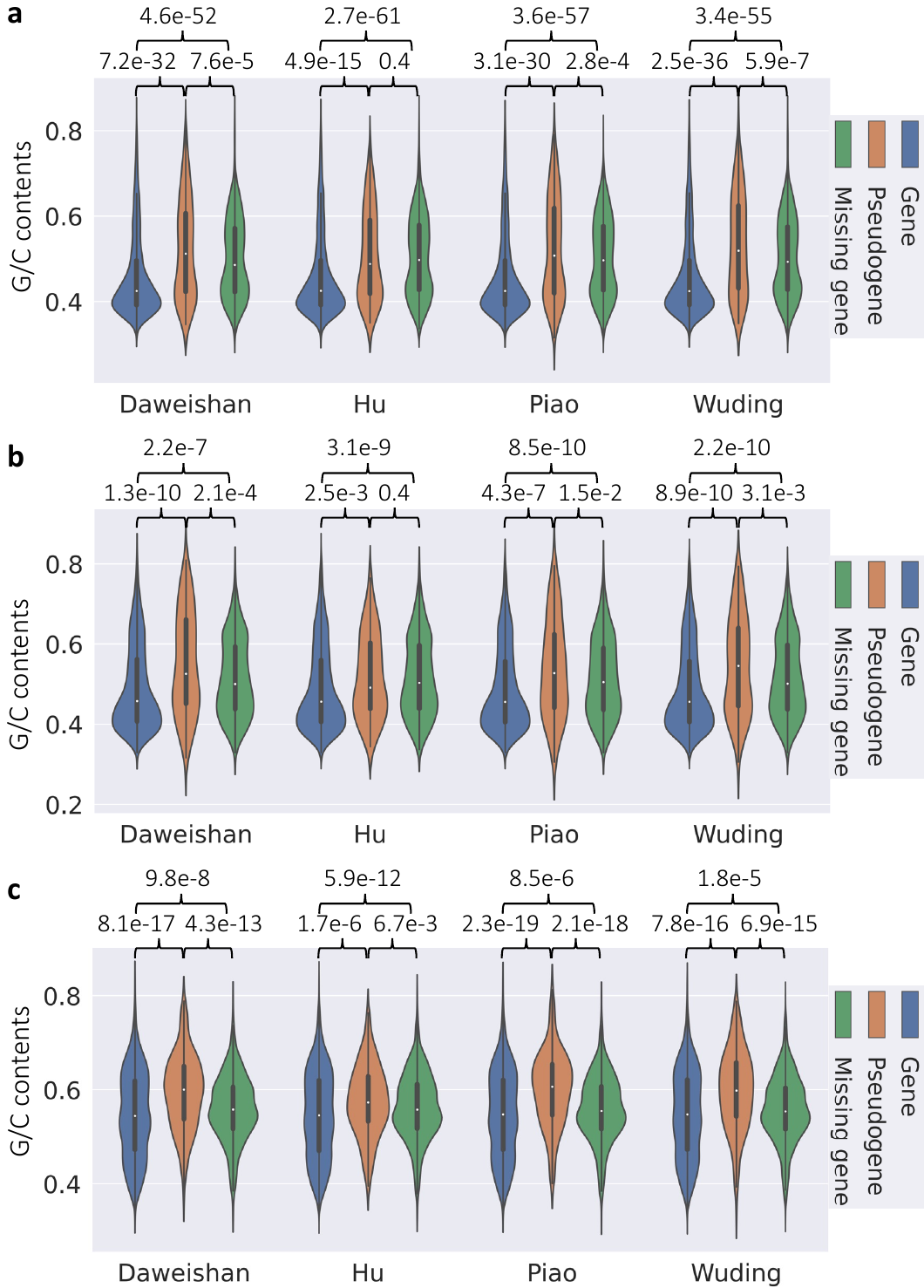

**Figure S4.** Comparison of G/C contents of true genes, pseudogenes and missing genes in the chicken genomes on different chromosomes. **a.** Comparison on macro-chromosomes (chr1-chr5 and chrZ). **b.** Comparison on intermediate-chromosomes (chr6-chr13 and chrW). **c.** Comparison on micro-chromosomes (chr14-chr39). Statistical tests were done using one-tailed t-test.

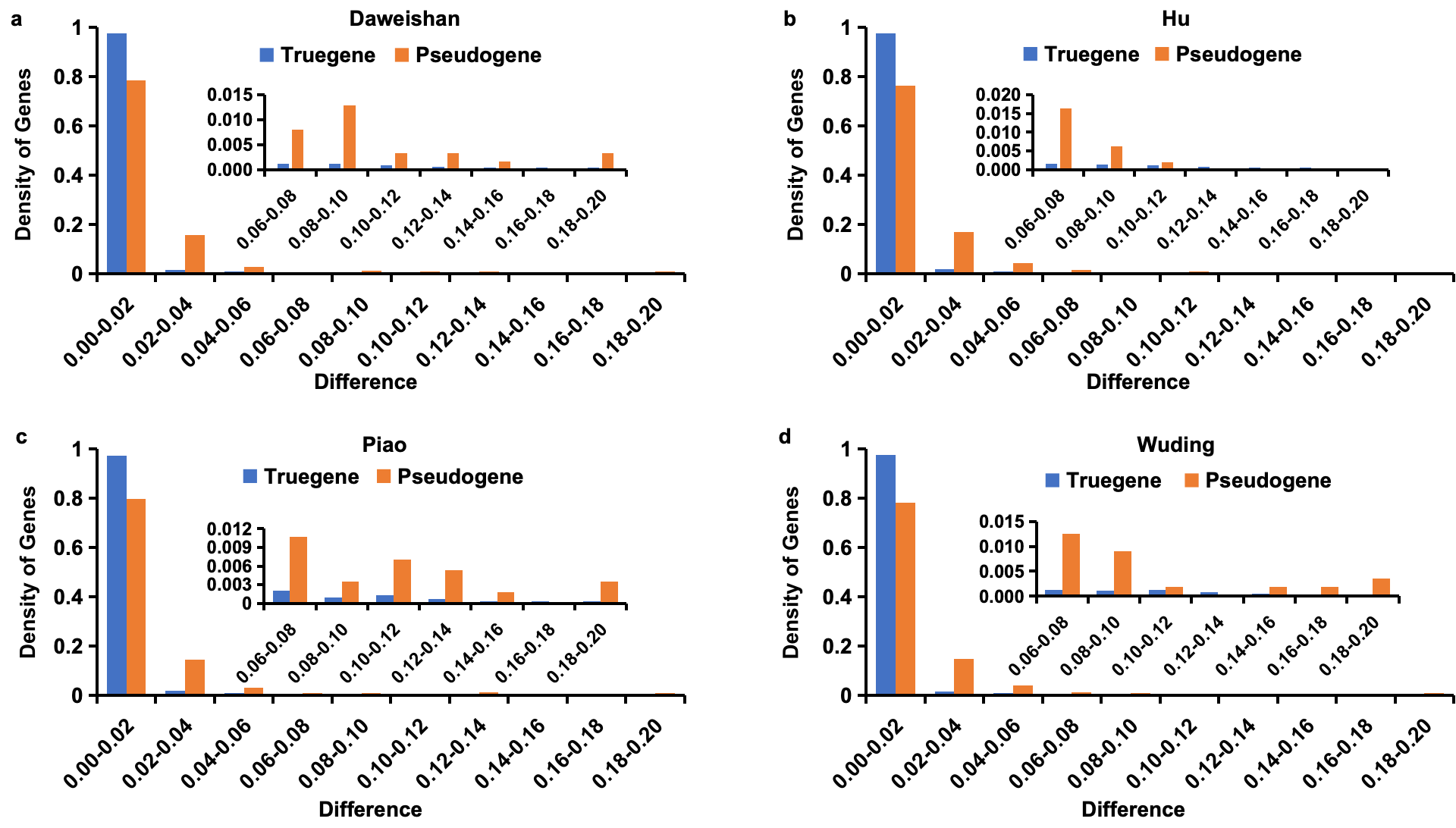

**Figure S5.** Distribution of difference rates of pseudogenes and true protein-coding genes in the four chicken breeds. The inset in each panel is the zooming-in view in the difference rates ranging from 0.06 to 0.2.
